# Autoregulation of a trimeric transporter involves the cytoplasmic domains of both adjacent subunits

**DOI:** 10.1101/2022.06.05.494857

**Authors:** Vanessa Leone, Richard T. Bradshaw, Caroline Koshy, Paul Suhwan Lee, Cristina Fenollar-Ferrer, Veronika Heinz, Christine Ziegler, Lucy R. Forrest

## Abstract

Membrane transporters mediate the passage of molecules across membranes and are essential for cellular function. While the transmembrane region of these proteins is responsible for substrate transport, often the cytoplasmic regions are required for modulating their activity. However, it can be difficult to obtain atomic-resolution descriptions of these autoregulatory domains by classical structural biology techniques if they lack a single, defined structure, as they may not be resolved or may be truncated or modified to facilitate crystallization. The betaine permease, BetP, a homotrimer, is a prominent and well-studied example of a membrane protein whose autoregulation depends on cytoplasmic N- and C-terminal segments. These domains sense and transduce changes in K^+^ concentration and in lipid bilayer properties caused by osmotic stress. However, structural data for these terminal domains is incomplete, which hinders a clear description of the molecular mechanism of autoregulation. Here we used µs-scale molecular simulations of the BetP trimer to compare reported conformations of the 45 amino-acid long C-terminal tails. The simulations provide support for the idea that the conformation derived from EM data represents a stable global orientation of the C-terminal segment under downregulating conditions. The simulations also allow a detailed molecular description of the C-terminal tail dynamics as well as its interactions with lipids, potassium ions, and the cytoplasmic surface of neighboring transporter subunits. Nevertheless, they do not provide information on the N-terminal segment, whose structure was not resolved by the structural studies. We therefore examined the possible interactions of the N-terminal tail by generating *de novo* models of its structure in the context of the EM-derived structure. The resultant full-length models of the BetP trimer provide a molecular framework for the arrangement of the terminal domains in the downregulated protein. In this framework, each C-terminal tail contacts the neighboring protomer in the clockwise direction (viewed from the cytoplasm), while the N-terminal tails only contact the protomer in the counterclockwise direction. This framework indicates an intricate interplay between the three protomers of BetP and, specifically, a multi-directionality that may facilitate autoregulation of betaine transport.

## Introduction

The ability of cells to move molecules across membranes is critical for accumulation of nutrients, extrusion of toxins, and responses to external stresses. Diverse transmembrane proteins dedicated to this process, called transporters, are abundant in all cells. A common stress factor for both human cells and bacteria is a change in osmotic conditions. Hyperosmotic stress, for example, presents a challenge for maintaining a constant cytoplasmic water activity and triggers cell shrinkage, oxidative stress, DNA damage, and apoptosis. One mechanism to counteract such an increase in external osmolality is to accumulate solutes such as betaine that are compatible with normal physiological functions. These “compatible solutes” can be synthesized by the cell, but synthesis requires regulation at the level of protein expression (1). The first response is therefore instead to take up compatible solutes from the environment, although this can prove a challenge because their concentrations are often at very low levels. To achieve the very high concentrations required to increase the total internal osmolality, uptake of compatible solutes must therefore be energized, for example, by coupling to an electrochemical gradient. However, the activity of compatible solute transporters must be tightly regulated such that the response only occurs under osmotic stress conditions. Regulation therefore must include, first, a sensor mechanism – to detect an associated stimulus such as cell turgor, membrane deformation, or internal ion concentration – and second, a mechanism to transduce this signal to the solute transport machinery. Among the many compatible solute uptake systems, one of the best studied is the glycine betaine transporter, BetP, from the soil bacterium *Corynebacterium glutamicum*. Impressively, BetP combines the osmosensor, the signal transducer, and the osmolyte transporter in a single macromolecule (2), and therefore provides an elegant prototype for transporter autoregulation in response to environmental change.

BetP is a homotrimer (3-5) in which the twelve transmembrane (TM) helices in the core of each protomer constitute the fundamental unit responsible for sodium-driven betaine uptake (6). In the absence of activation due to osmotic stress signals, this core domain mediates basal levels of transport (7, 8). The cytoplasmic N- and C-terminal tails function as osmosensors and osmoregulators (7, 9), in part by detecting high levels of the cytoplasmic concentration of monovalent cations, such as K^+^, to reach higher turnover rates. While the deletion of the ∼45-residue long C-terminal segment abolishes activation by osmotic stress, protein variants with N-terminal tail deletions retain the capacity for regulation, albeit requiring higher osmolality to achieve the same degree of activation (5, 7).

A key observation from structural studies of BetP is the abundance of interprotomer contacts that involve the C-terminal segments. In all available X-ray structures, the helical C-terminal segment of one protomer protrudes into the cytoplasm (Fig. 1a) and interacts with the cytoplasmic loops of an adjacent protomer (5), including with the segment called loop2, which connects transmembrane segments (TM) 2 and 3. Since TM3 undergoes some of the most consequential conformational changes during the transport cycle (10), the presence of this interaction implies crosstalk between the protomers during BetP regulation, as suggested previously based on low-resolution EM maps of 2D crystals (11). However, in the X-ray structures, one of the three C-terminal tails per trimer is also involved in forming the crystal lattice through interactions with the periplasmic surface of another trimer in the neighboring unit cell, a surface that is unavailable to the cytoplasmic C-terminal tails in cells. The presence of this plainly non-physiological interaction raises the possibility that this orientation of the C-terminal tail is a crystallographic artifact. More recently, a structure of BetP was determined using single-particle electron microscopy (EM) in the presence of amphipols to mimic the negative charges abundant in *C. glutamicum* membranes (12) unlike the dodecylmaltoside detergent micelles present in the crystals. In this EM structure, the C-terminal tail of each protomer also forms a long helix that interacts with a neighboring subunit, but it adopts a different orientation than in the crystals, lying parallel to the membrane plane along the outer rim of the protein (Fig. 1b). In this orientation, the C-terminal helix has the potential to interact more closely with lipids as well as with loop6, which connects TM6 to TM7, and with a different region of loop2. Adding to the enigma of why the C-terminal tail adopts different orientations, both crystal and EM structures were obtained under non-activating conditions (i.e., low K^+^ concentrations and osmolality).

**Figure 1.**
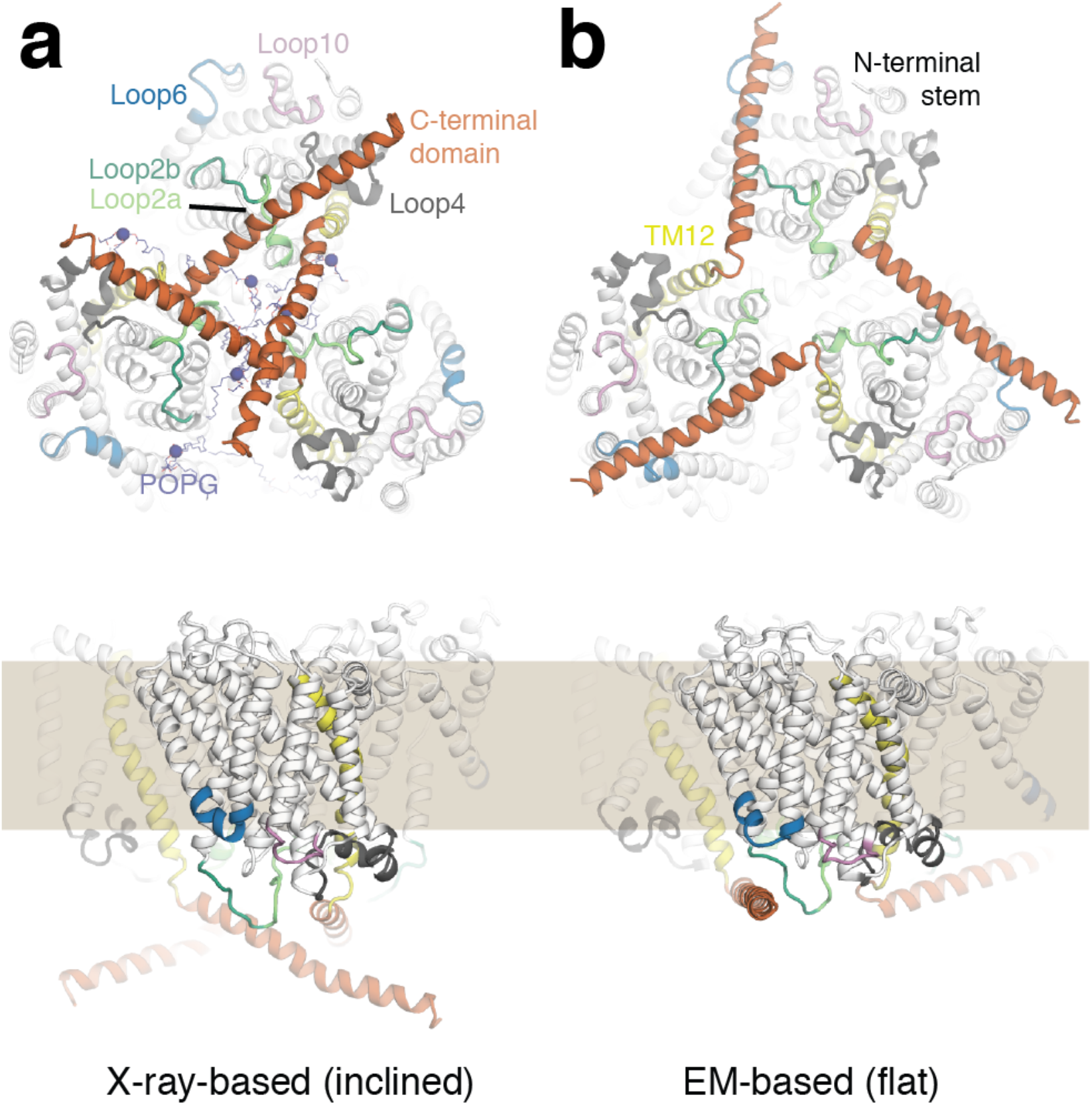
Arrangement of the C-terminal tails in the BetP trimer and position relative to the cytoplasmic loops. The relative positions of the C-terminal tails (dark orange) to the transporter domain and lipids differs in available structures of BetP. The protein is shown as cartoon helices and colored to highlight TM12 (yellow), loop2a (light green), loop2b (dark green), loop4 (gray), loop6 (blue) and loop10 (pink). POPG lipids resolved in the structure are shown in dark blue and as lines, with the phosphate atom as a sphere. Top panels show the view from the cytoplasm. Lower panels show the view from along the plane of the membrane (brown region) with one protomer in the forefront and the cytoplasmic regions at the bottom. **(a)** Structure derived from an X-ray crystal structure (PDB code 4C7R) with C-terminal domains inclined with respect to the membrane. **(b)** Structure derived from an EM map obtained in the presence of amphipols (12) with the C-terminal domains lying flat relative to the membrane surface.

Structural information relating to the ∼55-residue long N-terminal tail is significantly more limited. Secondary structure prediction methods suggest that this tail contains small segments of helix among regions that lack structural definition (9, 13). Accordingly, in the protein construct required for crystallographic studies, the first 29 amino acids were truncated and three glutamates (residues 44-46) were mutated to alanine to increase the helical content of this region (5). Despite aiding crystallization, only a few amino acids of this modified segment were resolved. The recent EM studies were performed with the wild-type sequence of BetP, but unfortunately the N-terminal segment was still not detected, suggesting that this portion is internally unstructured, or perhaps more likely, adopts an ensemble of different orientations with respect to the rest of the protein under the conditions used.

Looking beyond the protein to its environment, structural studies have also revealed densities for individual phosphatidylglycerol (PG) or cardiolipin (CL) lipid molecules that suggest specific contacts with the C-terminal helix and other protein regions, including loop2 (12, 14). Since the PG lipids were carried over despite detergent solubilization of the protein, it is possible that specific lipid-protein interactions are important for detecting and transducing osmotic stress signals to the transport domain of BetP.

A complete understanding of how the tails modulate the transporter domain cycling through its outward-open, occluded, and inward-open states, requires a molecular level description of the two cytoplasmic tails and their interplay with lipids, cations, and the rest of the protein. For the C-terminal tail, the most essential segment for regulation, it is not clear how the two observed conformations are related, nor how they behave in the absence of crystal contacts and in the presence of the lipid bilayer. In this work we investigated these two possible conformations (which we refer to as either inclined or flat, Fig. 1 and S1a), using µs-scale molecular dynamics (MD) simulations of the complete BetP trimer. Analysis of these simulations provides a detailed description of the conformational flexibility and interactions of the C-terminal tails, and the necessary context for predicting the structural features of the N-terminal tail. Using *ab initio* modeling we therefore construct models of full-length BetP, which indicate that autoregulation involves the cytoplasmic domains of both adjacent protomers.

## Materials and Methods

### MD simulations based on the crystal structure, with inclined C-terminal tails

To explore the crystal structure orientation of the C-terminal tails, the 2.7 Å-resolution symmetric inward-open BetP trimer (PDB code 4C7R (14), Uniprot code P54582) was selected. Side chain rotamers of asparagine, glutamine, and histidine residues were examined using the MolProbity webserver (November 2017) (15), and after visual inspection, seven of the 16 proposed were modified: Asn59(B), Gln215(C), Gln346(A), Gln439(B), Asn517(A), Asn517(C), where the letter in parentheses indicates the chain identifier. Histidine protonation was determined by inspecting possible hydrogen bond networks with other residues in the protein. All ionizable sidechains were treated with default ionization states, except for Glu161 and Asp439 (16). We retained the eight complete 1-palmitoyl-2-oleoyl-sn-glycero-3-phosphoglycerol (POPG) lipids present in the crystal structure and added water in protein cavities using DOWSER (17). The resulting molecular system was embedded in a hydrated POPG lipid bilayer, using GRId-based Force Field Input (GRIFFIN) (18). Sodium and chloride ions were added up to a concentration of 200 mM Na^+^, 100 mM K^+^, plus Na^+^ as counterions to neutralize the system. The protein and crystallographic lipids and ions were kept in place with harmonic restraints, while the lipid bilayer was equilibrated using a 20 ns-long MD simulation. The final bilayer size was ∼141 Å^2^. Only one of the C-terminal helices in the crystal structure is fully resolved, i.e., in protomer A. To extend the C-terminal tails of the other two protomers to the same length (residue 586), we aligned the C-terminal helix of protomer A onto the resolved regions of the other two protomers (residues 549 to 552 in protomer B and residues 549 to 561 in protomer C), and removed any overlapping water molecules. This system is shown in Fig. S1a and simulation lengths are provided in Table S1.

To assess the effect of potassium ions we prepared a second, identical system but with 200 mM Na^+^, 300 mM K^+^, and using Cl^−^ as counterions to neutralize the charges.

Both systems, i.e., with either 100 mM or 300 mM K^+^, were equilibrated by gradually releasing restraints on the protein over 23 ns: in the first 8 ns we applied harmonic restraints on the atom positions with gradually weaker force constants. In the next 5 ns we applied weak harmonic restraints (4 kcal/mol/Å^2^) on the root mean squared deviation (RMSD) of the backbone of each individual protomer, using the minimized structure as reference. We then applied harmonic restraints of 4 kcal/mol/Å^2^, and then 1 kcal/mol/Å^2^, during two additional 5 ns-long simulations, on the RMSD of all protomers simultaneously. In addition, restraints with the same force constants were applied to the ϕ and Ψ backbone dihedral angles in the C-terminal tails (residues 549 to 586), to retain their helical secondary structure. Finally, both systems were equilibrated without restraints for 100 ns, from which snapshots were extracted at 100 ns, 20 ns, 10 ns, and 50 ns and used as initial frames for Anton2 simulation replicates 1, 2, 3, and 4, respectively. The Anton2 simulation timescales were 4.8 µs (for run 1) or 2.4 µs (for runs 2-4), amounting to 12 µs for each crystal structure-derived, “inclined” system.

### MD simulations based on the EM data, with flat C-terminal tails

Due to the moderate resolution of the available EM structural data, which at the time of setup was ∼6.8 Å, we did not use the corresponding structural model as the initial configuration for our simulations. Instead, we placed the crystal structure components onto that structure. Specifically, each BetP protomer was superposed onto the EM structural model using the backbone atoms of the TM regions. The twelve TM regions are defined as residues 59-78, 82-124, 137-167, 178-207, 235-265, 277-294, 301-323, 326-348, 359-388, 394-426, 450-479, 488-510, and 515-544, respectively. The C-terminal helix from protomer A of the crystal structure was then fitted onto each of the C-terminal helices in the EM model. As above, the protein was embedded in a hydrated POPG bilayer (Fig. S1b) and two sets of systems were generated, with either 100 mM or 300 mM K^+^ in the aqueous solution. Equilibrations were carried out similarly to the crystal structure-derived, “inclined”, systems above, fixing the protein with harmonic restraints. However, an additional restraint was applied in order to keep the protein inside the EM map using MDFF (19), reducing the force constant gradually over 68 ns. While using MDFF we also applied restraints on the peptide bond dihedral angle and chirality, as well as backbone ϕ and Ψ dihedral angles to prevent any forces generated by MDFF from distorting these features. Finally, we equilibrated the system further for 100 ns without constraints; the Rosetta EM score (20) of the C-terminal tails was computed for this trajectory, to assess the agreement between each frame of the simulation with the experimental EM map.

Simulations were then carried out using Anton2 starting with either the last frame of the equilibration (run 1), or with one of four configurations taken from this trajectory, depending on their associated Rosetta EM score averaged over the three helices. Specifically, we selected the frame with the best agreement (EM score <–400 a.u. for run 2), worst agreement (EM score ∼–370 a.u. for run 3), and intermediate agreement (EM scores in the range –380-400 a.u. for runs 4 and 5) to the 6.8 Å EM map. Using these configurations, five Anton2 simulations were carried out for each K^+^ concentration (100 mM and 300 mM). The simulation time ranged from 2.4 µs (runs 2-4) to ∼7.9 µs (run 1 at 100 mM K^+^). The total simulation time for the systems with the flat C-terminal tail was ∼19.9 µs and ∼16.8 µs for the simulations with 100 mM K^+^ and 300 mM K^+^, respectively.

### MD simulation parameters

All minimization and equilibration steps prior to the Anton2 runs were performed with NAMD v2.12 (21). In all simulations, the CHARMM36 force field (22-24) was used to describe the protein, lipid, and ions, while TIP3P was used for the water molecules (25). The simulations were performed using a semi-isotropic NPT ensemble (T = 298 K and P = 1 atm, constant *x*:*y* ratio) and periodic boundary conditions were used in all directions. The time step was 2 fs.

Electrostatic interactions in the NAMD simulations were calculated using particle mesh Ewald (26) with a real-space cutoff of 12 Å. The cutoff for van der Waals interactions was also set to 12 Å, with a switching function starting at 10 Å applied to both electrostatic and van der Waals interactions. The Anton2 simulations used the (1,1,3)-RESPA multigrator (27) (the update interval for bonded, near, and far interactions, respectively), a 9-Å vdW cutoff, no long-range dispersion correction, and U-series electrostatics (28). The latter parameters were those selected by the Anton2 “chooser” code, which resulted in a pure hydrated POPG bilayer with similar properties to that obtained during parameterization of the CHARMM36 force field (SI section “Test of lipid dynamics on Anton2” and Fig. S2).

### MD simulation analysis

Several metrics were used to analyze the sampled conformations of the C-terminal tail during the simulations. The stability of the C-terminal tails was assessed by first aligning on backbone atoms of the TM regions (residue 59-78, 82-124, 137-149, 152-167, 178-207, 239-265, 277-294, 301-323, 359-388, 394-426, 450-479, 488-510, and 515-544) of the corresponding protomer and then measuring the RMSD of each C-terminal segment (residues 548 to 586). The secondary structure was computed using Definition of Secondary Structure of Proteins (DSSP) (29) using the MDtraj tool (30). The orientation of the C-terminal tail was measured using two angles: (1) a crossing angle between TM12 and the C-terminal helix and (2) a torsion angle defining the pivot of the C-terminal helix on the cytoplasmic surface of the trimer. The crossing angle was defined as the angle between two vectors connecting the centers of mass of the α-carbons of two sets of residues, namely residues 539-541 and 543-545 in TM12 and residues 551-553 and 562-564 in the C-terminal tail. Two pivot angles were measured, one that reflects the movement of the first half of the C-terminal tail and another that reflects the motion of the whole C-terminal tail. Both angles were defined by measuring the torsion angle between 4 points, each defined as the center of mass of the α-carbons of a group of residues. The first point is embedded in the core of the transporter domain, defined using residues 107-110 of TM2. The second point is the projection of point 1 along the axis of TM12, defined as a vector between the centers of mass of residues 532-535 and residues 540-543. The fourth point differs between the two torsion angles and is defined as either the center of mass of residues 554-557 or of residues 575-578, reflecting shorter or longer segments of the C-terminal tail, respectively. The third point is a projection of the fourth point onto the axis of TM12.

The Rosetta EM score (20) was used to measure the agreement between sampled configurations of the C-terminal tail and an independently-determined EM map of the symmetric inward-facing BetP resolved at 3.7 Å (12), which was obtained after the Anton2 simulations were completed and was not used during the setup of our simulations.

### De novo modeling of cytoplasmic N- and C-terminal tails

The N-terminal tail was modeled in the context of the BetP trimer complete with the C-terminal tail orientation identified in the simulations. To benchmark our modeling strategy, we also re-model the structure of the C-terminal tail within the same BetP trimer protein context. For all modeling we used the FloppyTail protocol (31) in Rosetta v3.9, which, in contrast to methods such as AlphaFold2 (32), was specifically developed for sampling the accessible conformational space of a portion of a protein. To avoid spending computational time sampling conformations in which the highly-charged terminal tails enter the hydrophobic core of the lipid bilayer, we created a slab of pseudo-atoms, by converting atoms P, C13, C2, C23, C27, C211, C215, C32, C36, C310, and C314 of the lipids in the cytoplasmic leaflet of the POPG bilayer. A repulsion term between the newly-modeled tail and these pseudo-atoms was added to discourage exploration of this space. The three-residue long fragments used to sample the backbone rotamers of the terminal segments were obtained using the ROBETTA server (http://robetta.bakerlab.org/fragmentsubmit.jsp). We excluded sequences homologous to BetP, to prevent fragments from X-ray structures of BetP from being used. For the C-terminal tail benchmark, this choice avoids bias toward the known structure. For the N-terminal tail models this choice prevents inclusion of the structures obtained for the crystal construct, which is a modified version with a higher propensity to form helical structures (EEE44/45/46AAA).

Several sets of 100,000 cytoplasmic tail models were generated in the context of the trimer. Instead of identifying a single structure as the true terminal tail structure, global metrics were used to assess plausible secondary structure (with DSSP) and interactions with the rest of the trimer. To assess the spread of the sampled models we clustered them in structural space using Calibur (33), allowing the protocol to select the optimal clustering threshold. To assess the correlation between the interaction of the N-terminal with the C-terminal tail and the internal interactions of the N-terminal tail, two coordination numbers (*C*) were defined:

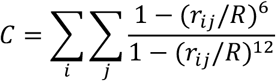

where *i* and *j* refer to each non-hydrogen atom in each residue pair considered, *r*_*ij*_ is the distance between these atoms in the given model and *R* is 5 Å. The more contacts formed between the two groups of atoms, the larger the value of the coordination number. We considered two sets of interactions with this metric. The first analysis measured the contacts between residues 548-586 and 1-55, i.e., between the C- and N-terminal tails. The second analysis measured the interactions between two helical portions (residues 10-31 and 36-49) in the N-terminal tail. The coordination numbers were computed using the Collective Variable (COLVAR) Dashboard plugin in VMD (34). The models were clustered based on the two-dimensional space defined by these two coordination numbers using the METAGUI analysis interface (35, 36).

## Results

The primary question that arises from comparing the structures of BetP is the preferred orientation of the C-terminal helix under non-activating conditions (low osmolality), i.e., whether this segment is more likely to be inclined with respect to the surface of the membrane, as in the crystal structures, or to lie flat alongside the surface of the protein as in the cryo-EM data. To address this question, we carried out µs-scale MD simulations of BetP in a hydrated POPG lipid bilayer, to examine the behavior of the two alternative orientations, starting with the inclined orientation.

### Simulations of the inclined C-terminal conformation derived from a crystal structure

#### The C-terminal helix does not remain inclined in the absence of crystal contacts

The inclined orientation of the C-terminal helix is observed in all reported crystal structures, which include many different conformations of the membrane-spanning transport domain. Of these conformations, the inward-open state is the preferred conformation in resting conditions in the presence of saturating amounts of sodium but in the absence of betaine substrate (16). Therefore, here we simulated the trimer with all three transport domains in inward-open conformations (PDB code 4C7R), and with all three C-terminal helices in an inclined orientation. In the initial simulations, the aqueous solution contained a sub-activating (100 mM) K^+^ concentration. For these simulations, we assessed the structural drift of the C-terminal segment and the persistence of the helix. The initial X-ray-derived conformation was lost after a few hundred nanoseconds, due to changes not only in the orientation of the helix relative to the membrane domain, as measured using the backbone RMSD of the tail (Fig. 2a), but also due to changes in helicity (Fig. 2c). The extended orientation was not revisited during a 4.8 µs-long simulation. To evaluate whether this instability of the inclined conformation reflected the choice of initial structure, we performed three additional 2.4 µs-long simulations starting from different frames extracted from the equilibration stages. Again, in all the simulations, at least one of the three C-terminal segments diverged rapidly from the inclined structure (Fig. S3a-d) and its helicity was lost around residue Arg565 (Fig. S4a-d). Notably, the helix involved differed; in the first simulation, the broken C-terminal helix was in protomer A (i.e., the one that was fully resolved in the crystal structure); and in the third run, a similar breakage occurred in protomers B and C (Fig. S4e).

**Figure 2.**
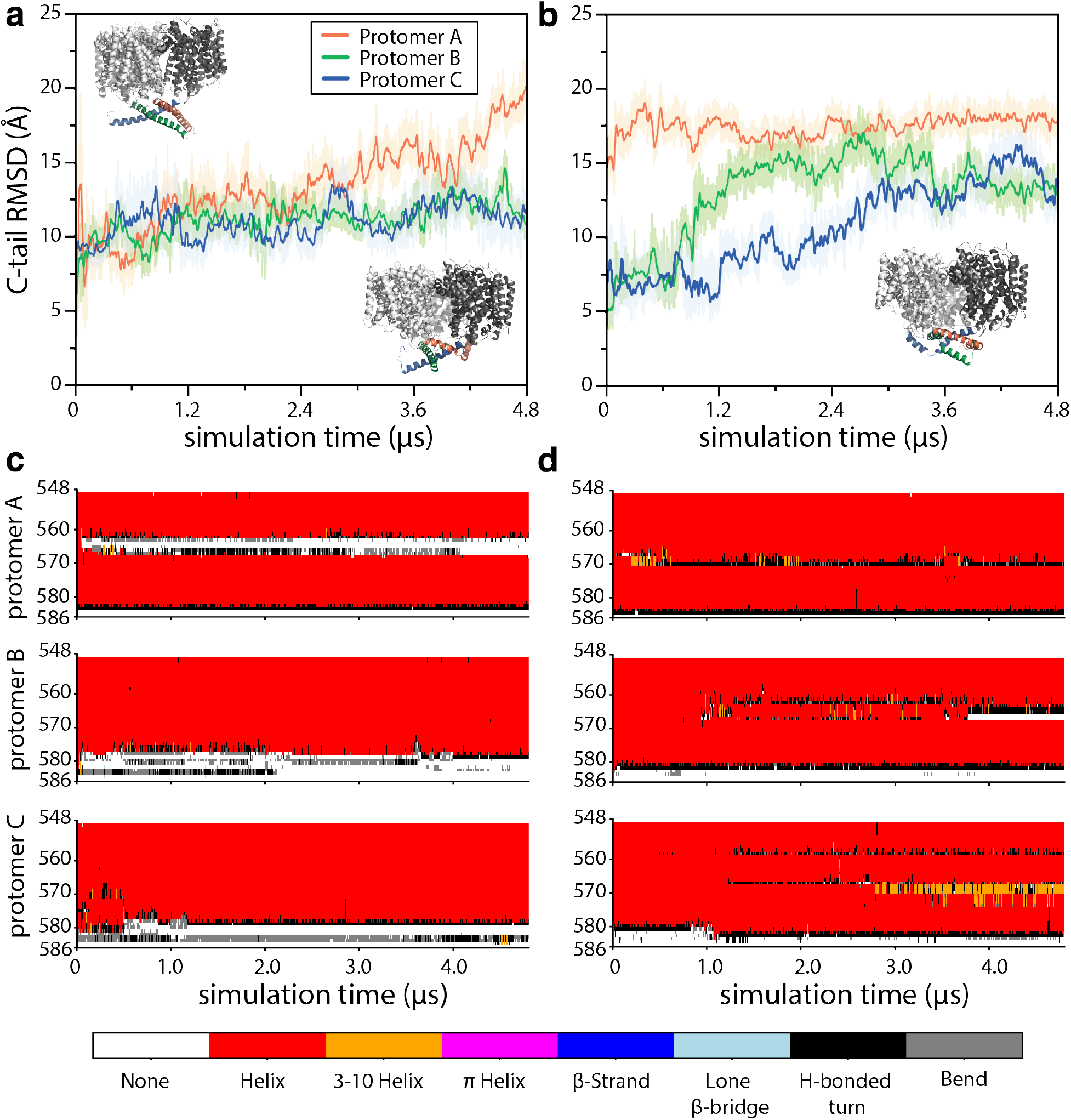
Structural stability of the C-terminal tails in the conformation derived from the crystal structure (all-inclined). **(a, b)** RMSD of each of the three C-terminal tails in BetP after aligning the trimer using the transmembrane regions (as defined in the methods section). The reference structure is the initial structure as shown inset in panel a (top left), while the structure of the final frame in each simulation is shown in the bottom right of the corresponding panel. **(c, d)** The secondary structure as a function of simulation time computed using DSSP and plotted for every residue in each C-terminal tail using the indicated color scheme. Results are shown for the two longest simulations with 100 mM K^+^ (panels a and c) and 300 mM K^+^ (panels b and d) and are compared with data from repeated trajectories in Figures S3 and S4.

#### Potassium ions do not stabilize the C-terminal helix in the inclined orientation

The K^+^ concentration used in the abovementioned simulations was below that required for full activation (K_m_ ∼220 mM). Given the role of the C-terminal tail in regulation, we considered the possibility that the inclined orientation might require high concentrations of K^+^ to be stabilized. Hence, we simulated the trimer with the inclined helices in the presence of 300 mM K^+^. Again, in each of four repeat simulations of 2.4 to 4.8 µs length, one or more of the C-terminal helices deviated from the inclined orientation (Fig. 2b and Fig. S3e-h) and irreversibly lost helicity (Fig. 2d), suggesting that a high K^+^ concentration is not sufficient to stabilize this orientation.

#### The C-terminal region not involved in crystal contacts samples interactions observed in the crystal structure

The crystal lattice contacts formed by the extended C-terminal helix involve residues toward the end of the C-terminal segment, after the position at which breaking of the helix was typically observed in our simulations. Here, we defined the hinge point to be around residue Ala564 since the A564P substitution abolishes regulation by BetP (37) (Fig. 2c-d and Fig. S3). However, the segment prior to the breakage point (residues Val548 to Ala564) also contains significant secondary structure, and thus, it is plausible that the conformation of the first half of the C-terminal tail is stable and physiologically relevant. Indeed, during our simulations this first segment of the C-terminal tail did not deviate substantially from the initial conformation derived from the X-ray structure (Fig. S5a-d).

To understand what defines the orientation of the first helical segment, we computed its interactions with the rest of the protein in the simulations. Several intraprotomer contacts found in the crystal structure (Table S2) were also sampled in the simulations (Table S3, Fig. S6a,c,d). These contacts involve sequence-adjacent residues from the end of TM12 (e.g., Leu544, Ser545, and Asn546 interacted frequently with Tyr550 or Leu551) as well as residues in loop2a (e.g., Lys121, Phe122 and Ile125 interacted frequently with Ile549, Tyr550 or Tyr553) and loop4 (Tyr206 and Arg210 with Val548 and Ile549). The interactions with loop2a and loop4 are possible because the first few residues of the C-terminal tail are positioned toward the center of its own protomer rather than at the lipid-exposed periphery (Fig. 1a).

In addition to the observed intraprotomer contacts, this first segment of the C-terminal tail interacted with both adjacent protomers due to the crisscrossing of the three tails (Table S2 and S3, Fig. 1b, Fig. S6a,d). In the clockwise direction (viewed from the cytoplasm), contacts with Asp131 in loop2b were frequently observed in the simulations and involved Arg558 and Arg562. The remaining contacts involved the C-terminal tails from either the clockwise (with residues 549 to 556) or counterclockwise protomers (with residues 560 to 578), the most frequently observed being Glu552 with either Arg567 or Arg568.

Taken together, the RMSD and contact analysis indicate that the first and second segments of the C-terminal tail (residues 548-564 and 565-590) in the C-terminal tail should be considered separately. While the segment following Ala564 is highly dynamic during the simulations, the segment prior is substantially less mobile and samples intra- and inter-protomer contacts that are also observed in the experimental structure. These findings suggest that the portion of the C-terminal tail not involved in lattice contacts may in fact be adopting a physiologically-relevant orientation in the X-ray structure, one that may be supported by frequently-formed interactions with the cytoplasmic loops in the same protomer.

#### The conformation of the first part of the C-terminal tail does not change with the addition of potassium ions

Although 300 mM K^+^ did not stabilize the entire length of the C-terminal tail (Fig. 2), it is also possible that K^+^ plays a more localized structural role, for example, in modulating the interactions between the first portion of the C-terminal tail and the cytoplasmic surface of the BetP trimer. To examine this possibility, we analyzed the C-terminal tail RMSD and contacts in simulations performed with 300 mM K^+^. The RMSD of residues 548-564 indicates a relatively stable conformation (Fig. S5e-h), similar to that observed in the 100 mM K^+^ simulations. In addition, the contacts between this C-terminal region and the rest of the trimer involve the same residues independent of the concentration of K^+^ (Table S3, Fig. S6b). Furthermore, the K^+^ ions themselves form essentially no contacts with the C-terminal tail (Fig. S7a-b), even at a high concentration of K^+^, and therefore do not compete directly with the interactions between the C-terminal tail and the rest of the trimer.

#### Cations interact frequently with Asp470 in TM10

One of the activation stimuli for BetP (K_m_ ∼ 220 mM) is the presence of a high concentration of K^+^ in the cytoplasm (8). Since K^+^ ions were not observed to bind to the C-terminal tail in our simulations, we analyzed the trajectories for other regions in the trimer that consistently form contacts with K^+^ ions (Fig. S7a, S7b). A few cytoplasmic residues stand out as frequently forming contacts in all three protomers, especially Asp470 from TM10 and Ser293 from TM6, for an average of 21% and 22% of the 300 mM K^+^ simulations, respectively (Fig. S7b). These two residues contribute to a crevice between TM10 and the flexible loop connecting TM6 and TM7 (loop6) (Fig. 3a, 3e). K^+^ ions accumulated in this crevice more than in any other region on the cytoplasm face of the protein and the occupancy increased four-fold with the K^+^ concentration (Fig. 3b).

**Figure 3.**
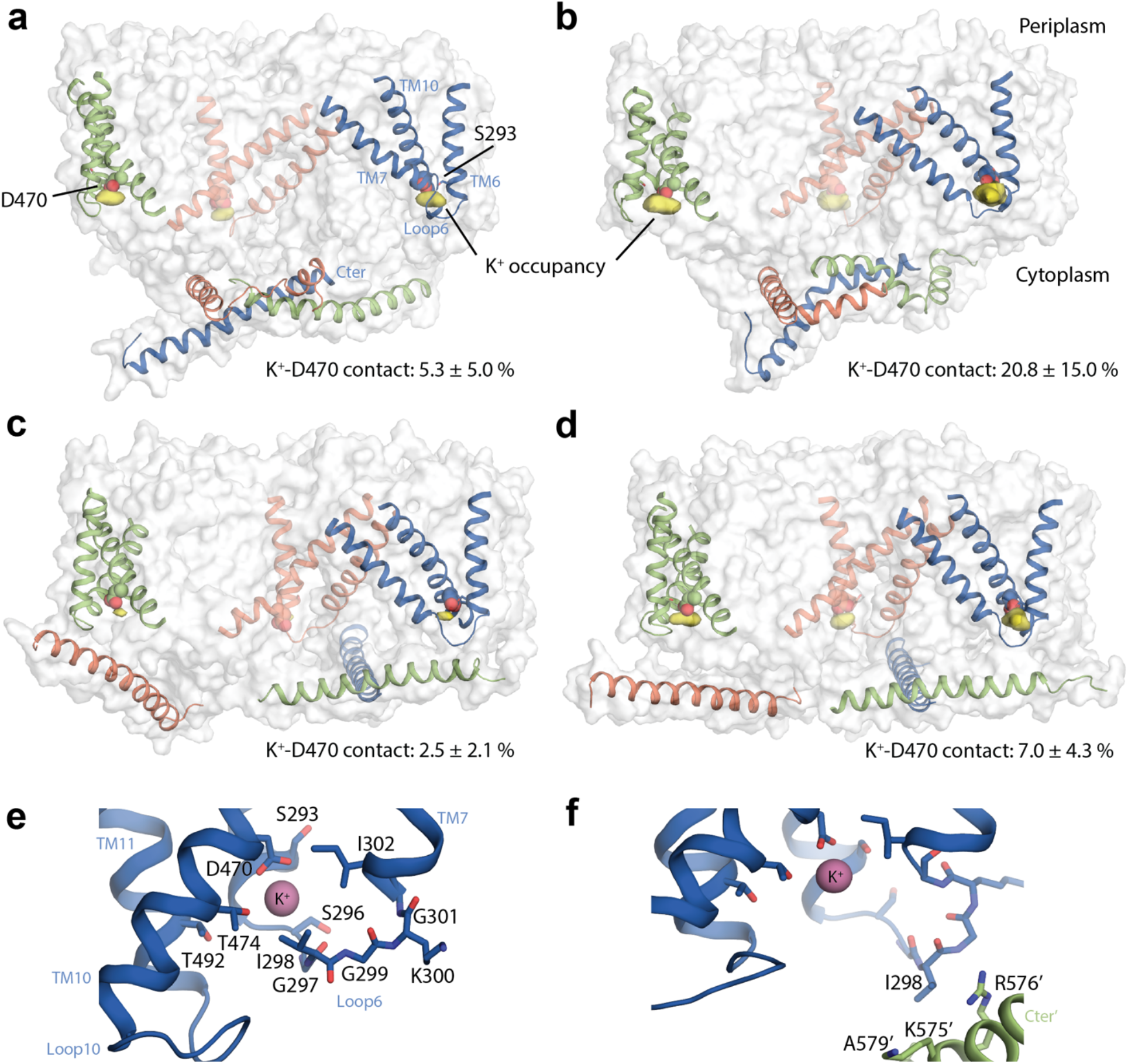
Cation interaction site predicted by MD simulations of the BetP trimer. **(a-d)** Potassium ion occupancy map (yellow surface), showing only densities close to the cytoplasmic surface and present in all three protomers, for clarity. Ion-protein contacts are defined as any K^+^ ion within 3.5 Å of any non-hydrogen atom of residue Asp470. The protein is shown in gray surface representation and as cartoon helices for TM6, loop6, TM7, TM10 and the C-terminal tail, colored by protomer (orange, blue and green). Asp470 is shown as spheres, and Ser293 is shown as sticks. Simulations were initiated with either **(a, b)** the inclined orientation of the C-terminal tails found in X-ray crystallographic structures, or **(c, d)** the flat orientation observed in EM maps. The bulk concentration of K^+^ was either 100 mM (a, c) or 300 mM (b, d). **(e, f)** Closeup of the potassium ion when coordinated by Asp470 in simulations in which the C-terminal tail is inclined (e), or in a flat orientation (f). Representative snapshots with high coordination numbers are shown with the ion (sphere), and nearby amino acids (sticks) from TM6, loop6, TM7, TM10, loop10 and TM11 (cartoon helices). In (e), the C-terminal tail of the adjacent protomer contacts residues in loop6. Individual residues are labelled. Segments from an adjacent protomer are marked with prime (*‘*).

Given the observed high accessibility and moderate occupancy of the Asp470 crevice, we considered the possibility that this site can non-specifically bind all monovalent cations. The aqueous solution used in the simulations contained 200 mM Na^+^ (required to counteract the negative charges of the POPG lipids) and so we also analyzed the occupancy of Na^+^ ions in this region. At this concentration, Na^+^ ions also interact frequently with Asp470 (Fig. S7a-d, g), albeit to a lesser extent than K^+^ ions, for 5.4% of the simulations with 100 mM K^+^ (Fig. S7c). Ser293 binds Na^+^ ions, on average, 7% of those simulations.

### Simulations of the flat C-terminal conformation derived from cryo-EM data

Recently-determined EM structures suggest that, in the absence of K^+^ and high osmolality, i.e. under non-activating conditions, the C-terminal tail forms a long, relatively-straight helix that lies flat along the trimer cytoplasmic surface (Fig. S1)(12). Importantly, the EM structure lacks the issues associated with the crystal contacts discussed in the previous section, although its resolution hinders clear identification of many of the specific interactions that favor the flattened orientation of the helix, including those with the lipids. Moreover, the EM data was obtained for BetP trimers in amphipols, which mimic the high density of negative charges present in the *C. glutamicum* membrane but raise the question of whether the presence of a lipid bilayer might lead to a different orientation of the C-terminal tail. Thus, here, we evaluate the stability of this flat conformation in microscopic detail and in the presence of a lipid bilayer. Specifically, we used information from a 6.8 Å cryo-EM map to construct the trimer orientation by first fitting the longest X-ray-resolved C-terminal helix (protomer A) into each C-terminal segment in the EM map and then restraining the protein into the map using MDFF (19) during equilibration (see Methods). As with the inclined helical conformation, unconstrained simulations were performed with both low K^+^ (100 mM) and high K^+^ (300 mM) ionic environments, but were up to 7.9 µs in length.

#### The EM-derived conformation remains stable

In the simulations of BetP carried out with the lower concentration of K^+^, the C-terminal helix in the EM-derived conformation did not deviate greatly from the initial flat structure, even over the course of a 7.9 µs-long simulation (Fig. 4a, Fig. S8a). The RMSD values for the C-terminal tail flatten off at ∼4-7 Å, in stark contrast to the rapid and large (15-20 Å RMSD) structural changes observed in the simulations of the X-ray-derived state of BetP (Fig. 2a, Fig. S3), suggesting that the C-terminal helix retains a flat conformation. Notably, the helicity was preserved throughout this long simulation, aside from the last seven residues (580-586, Fig. 4c), which are also poorly resolved in the EM map. These observations held true for four repeat simulations, where the global C-terminal conformation (Fig. S8b-e) and helical content (Fig. S9a-e) remained relatively stable.

**Figure 4.**
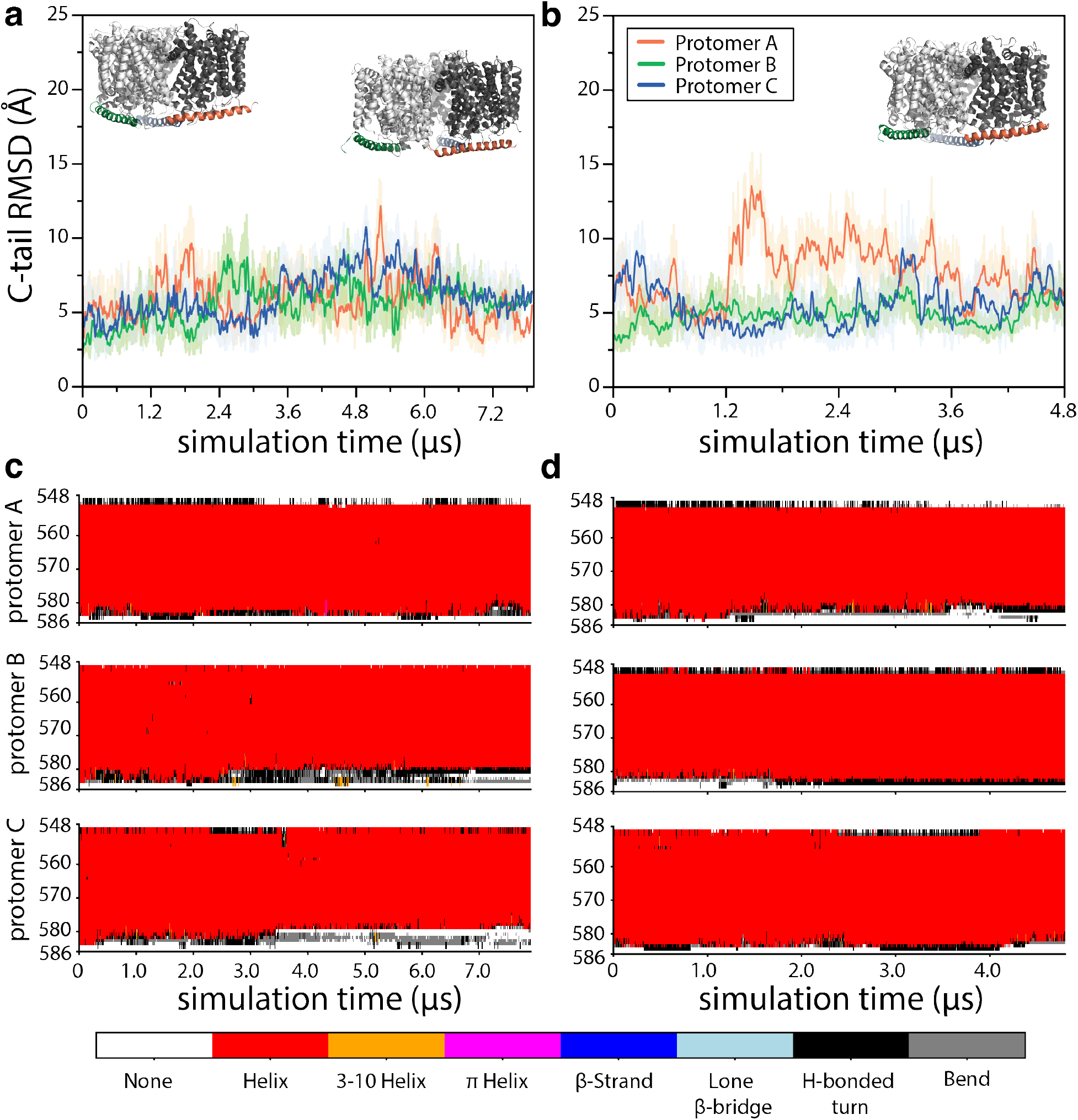
Structural stability of the C-terminal tails in the conformation derived from the EM structure (flat). **(a, b)** RMSD of each of the three C-terminal tails in BetP after aligning the trimer on the transmembrane regions. The reference structure is the initial model, as shown inset in panel a (top left), while the structure of the final frame in each simulation is shown in the top right of the corresponding panel. **(c, d)** The secondary structure computed using DSSP, plotted for every residue in each C-terminal tail as a function of simulation time, using the indicated color scheme. Results are shown for the two longest simulations with 100 mM K^+^ (panels a and c) and 300 mM K^+^ (panels b and d) and are compared with RMSDs in repeated trajectories in Figure S9.

These RMSD values were computed with respect to the structural model built based on 6.8 Å-resolution EM data, which was used as the starting point for the simulations. To relate the observed fluctuations during the simulations more directly to an experimental observable, we used a metric of the overlap between the EM map and a computed map from each simulation frame, namely the Rosetta EM score (20) (see Methods). Since an independently-obtained, higher-resolution EM map (3.7 Å) was available at this point, this new map was used for analyzing the simulations(12). According to this metric, most of the simulation snapshots overlap with the experimental map to a similar degree (Fig. 5) but can fluctuate around the mean value (Fig. S10). Figures 5c and 5d provide a structural representation of the fit between a simulation snapshot and the experimental data, illustrating how the C-terminal helix maintains a flat configuration during the simulations while accessing various alternative orientations around that position that all have a similar degree of overlap with the experimental map.

**Figure 5.**
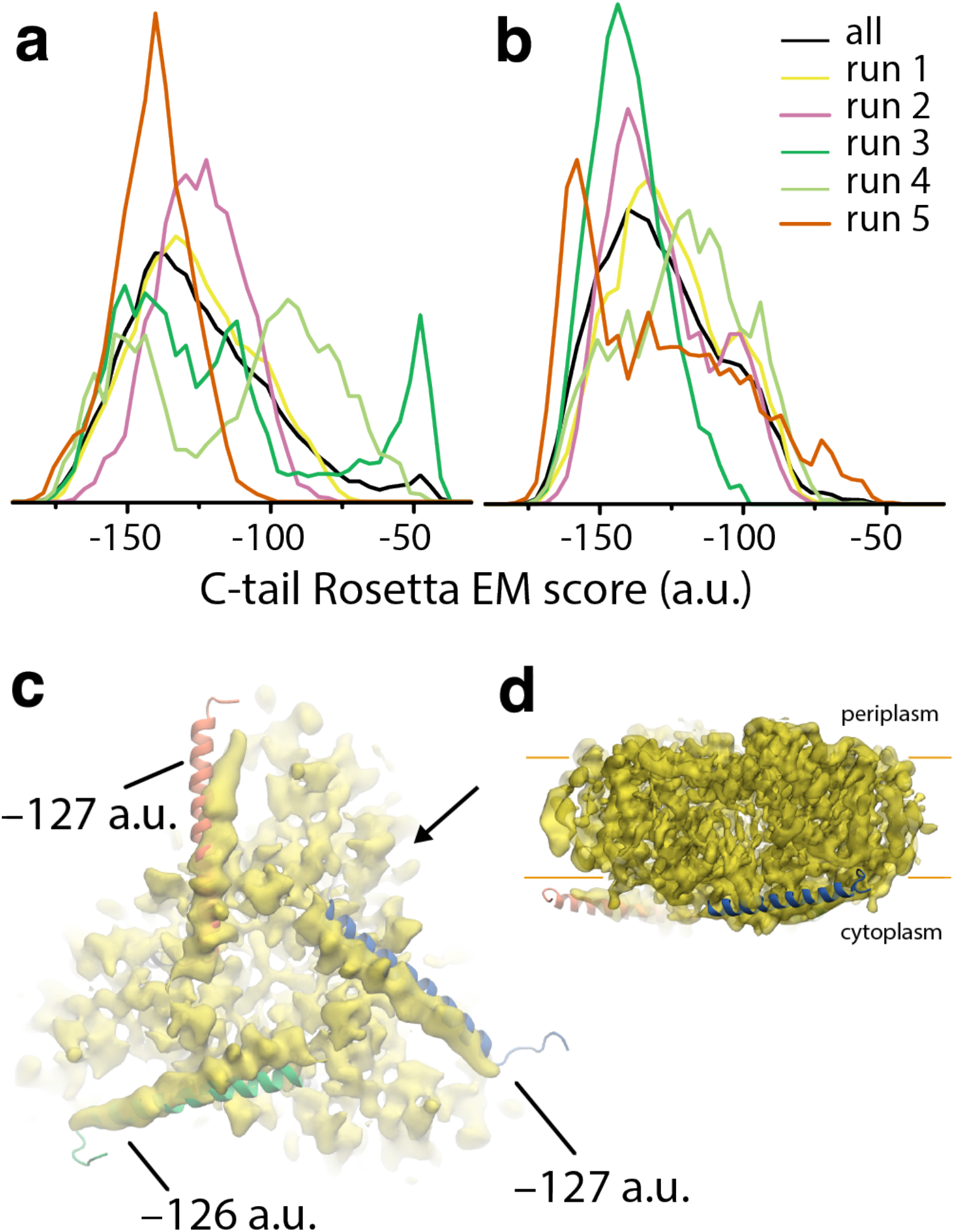
Agreement of the flat C-terminal tail conformations in simulations of the BetP trimer with EM maps. **(a, b)** The computed Rosetta Electron Microscopy (20) score, computed for each of the three C-terminal tails in each independent simulation (colored lines), or summed over each set of simulations (black lines), initiated with EM-derived structures, is shown as a distribution over the simulations carried out with either 100 mM K^+^ (a) or 300mM K^+^ (b). The accumulated simulation time is > 52.2 µs and 43.2 µs, respectively. **(c**,**d)** Simulation snapshot, shown as cartoon helices, compared with the EM map (yellow), and viewed from **(c)** the cytoplasm, or **(d)** the plane of the membrane, along the direction marked with an arrow in panel c. The Rosetta EM score of all three C-terminal tails is near the median value (∼–127 a.u.). Fluctuations in the C-terminal helix positioning can occur along its full length (blue), at the end of the C-terminal tail (orange), or close to the connection with TM12 (green). In (d), the approximate boundaries of the membrane are marked with orange lines.

#### The C-terminal helix derived from the EM maps remains parallel to the membrane, but pivots on the trimer surface

The abovementioned structural analyses indicate that the C-terminal helix of BetP does not drift away from its position in the EM map, but rather that it can take substantial excursions, or fluctuations around that average position during the simulations. To describe these variations in C-terminal tail orientation we developed three different geometrical metrics. First, to assess whether if the helix truly remains flat, i.e., parallel to the membrane plane, we measured the crossing angle between a vector describing TM12 and another vector describing the C-terminal segment (up to Ala564, i.e., the breakage-point in the inclined simulations; see Methods). In the simulations based on the EM data with the lower concentration of K^+^, the value of this angle remains around 70-110º (Fig. 6c), substantially smaller than that obtained for simulations based on the crystal structure (110-150º; Fig. 6a), indicating that the C-terminal helix indeed remains flat in the former simulations.

**Figure 6.**
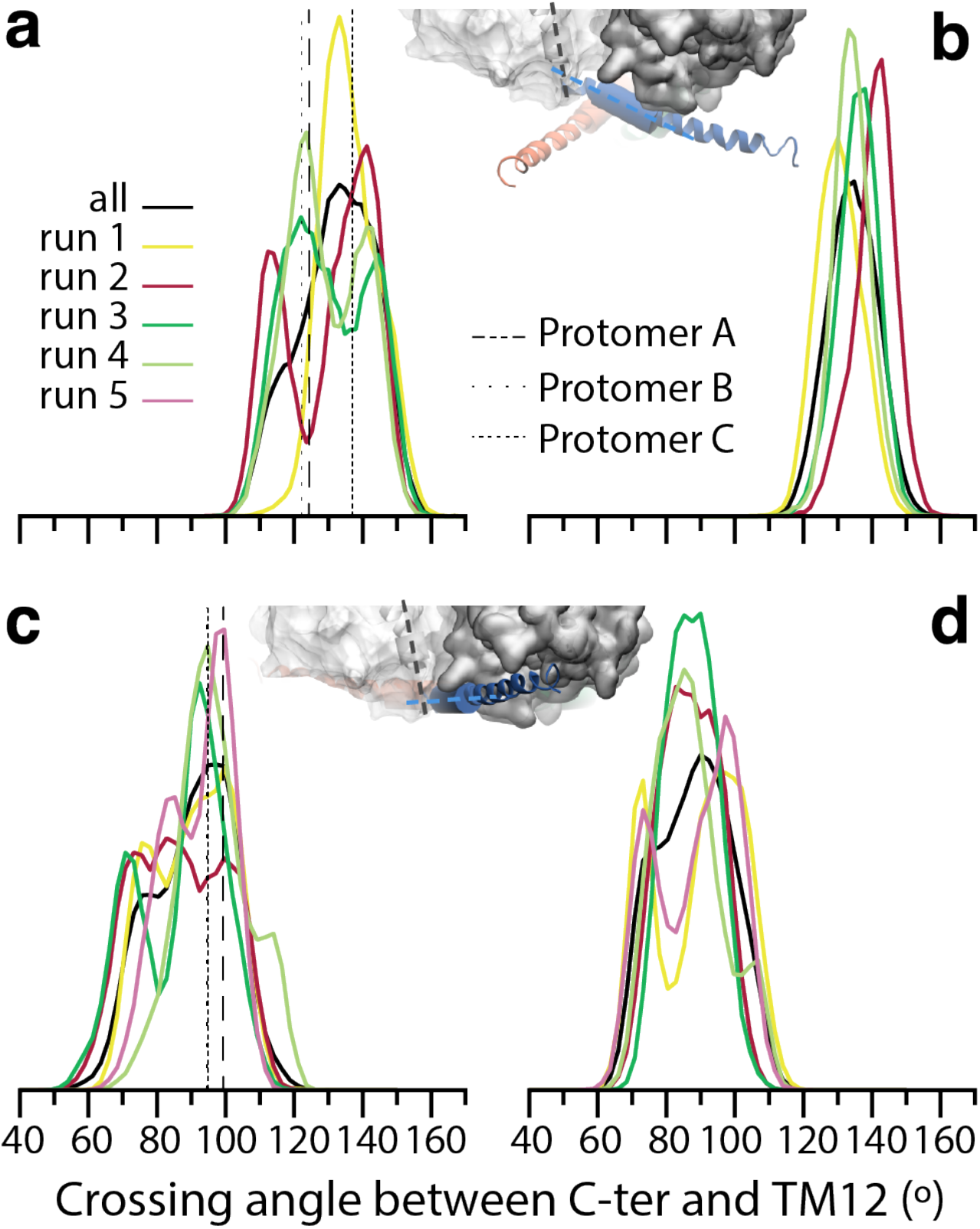
Inclination of the C-terminal tail of BetP in simulations derived from crystallographic (inclined) and EM data (flat). The crossing angle between TM12 and the C-terminal helices is shown as a distribution over each simulation, combining the sampling of the three protomers (colored lines), or summed over all trajectories (solid black lines). The crossing angle was defined using vectors for TM12 and the C-terminal tail that connecting the center of mass of the Cα atoms in two helical segments (539-541 and 543-545 for TM12; 551-553 and 562-564 for the C-terminal tail). Data are shown for simulations of **(a, b)** the crystal structure with either 100 mM K^+^ (a) or 300mM K^+^ (b) and **(c, d)** of the EM structure, with either 100mM K^+^ (c) or 300mM K^+^ (d). The values of the crossing angles in the corresponding crystal and EM structures are indicated using vertical black dotted and dashed lines in panels a and c. Inset structures indicate the segments in TM12 (gray cartoons) and the C-terminal tail (colored cartoons) used to define the crossing angle (cylinders and dashed lines), and their orientation relative to the transmembrane segments, shown in surface representation.

Having established that the C-terminal helix of BetP remains flat in these simulations, we analyzed the extent of any pivoting motions on the surface of the trimer. Such movements dictate whether the C-terminal tail can interact with the transporter domain loops or with the surrounding lipids. Accordingly, we defined a pivot angle using TM12 as a reference, such that values < 90° reflect movements of the C-terminal toward the protein surface and values > 90° represent fluctuations toward the POPG bilayer (see Methods). When measured over the entire length of the C-terminal segment, the distribution of the pivot angle is narrow, and centered around the values obtained for the initial structure (∼90º, Fig. 7a). Representing these vectors as lines illustrates the degree of fluctuation that can occur over the length of the C-terminal tail (see inset to Fig. 7a), which is likely to correspond to variation in the C-terminal tail contacts with the neighboring (clockwise) protomer or the lipid headgroups.

**Figure 7.**
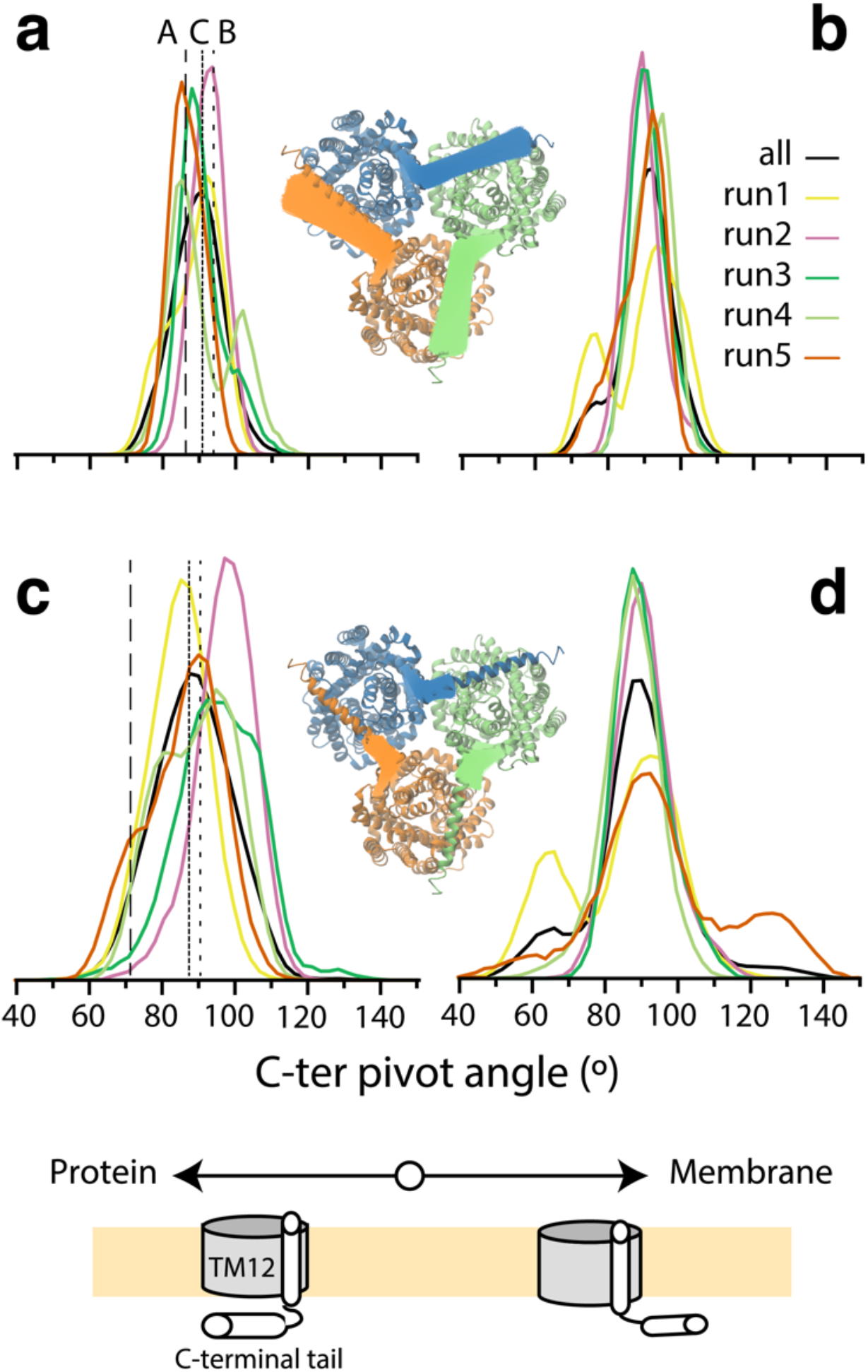
Pivoting of the C-terminal helix of BetP in simulations derived from the EM data (all-flat). The orientation of the C-terminal helix is plotted as a distribution over each simulation trajectory (colored lines) and averaged over all five runs (solid black lines). **(a, b)** Torsion angle defined using the full-length of the helical segment of the C-terminal tail, with the endpoint at the center of residues 575-578). **(c, d)** Torsion angle defined for the first helical segment of the C-terminal tail, with the endpoint at the center of residues 554-557. Simulations were carried out with either 100 mM K^+^ (a, c) or 300 mM K^+^ (b, d). The angles obtained for the corresponding experimental structure are shown as vertical black lines in panels a and c. Insets illustrate the position of the vectors defining the pivot angles for each C-terminal tail (green, orange, and blue lines), in 32909 frames from the 7.9 µs-long trajectory carried out in the presence of 100 mM K^+^. The protein is viewed from the cytoplasm. Schematics illustrate how the orientation of the C-terminal tail towards either the lipid bilayer or the center of the neighboring protomer is reflected by larger or smaller pivot angles, respectively.

Interestingly, as proposed previously (9), we also noticed a disconnect between movements in the first and second halves of the C-terminal tail. Hence, we defined a shorter version of this angle to separate out the behavior of these two segments. With this metric, the distribution is broader, and the initial structures have different values depending on the subunit (Fig. 7c), indicating local variations in the segment of the C-terminal tail close to TM12 (see green helix in Fig. 5c and inset to Fig. 7c). Such repositioning may be facilitated by changes in helicity at the very beginning of the C-terminal tail (residues 548-550; Fig. 3c, Fig. S9a-e) that affect the length of the linker or simply by repositioning of the helix with respect to TM12 (e.g., Fig 2c, orange C-terminal helix; Fig. S6c), even while maintaining contacts at the other end of the domain. Together, these analyses illustrate that, although the C-terminal segment is primarily helical, it exhibits complex behaviors beyond those that might be imagined for a rod-like structure.

#### The C-terminal helix in the flat orientation does not contact the counterclockwise protomer

It is notable, given the presence of a bilayer comprised entirely of negatively charged lipids (POPG in these simulations), that the highly positively charged C-terminal helix does not simply swing outwards to interact with those negatively charged headgroups (Fig. 7). Such a reorientation is presumably less favorable than existing interactions between the transport domain and the C-terminal tail, analogous to those observed in the inclined conformation (Table S3, Fig. S6). Indeed, we observe that the C-terminal tail forms several persistent contacts involving the same residues in TM12. However, there are also interactions with TM12 that are unique to the flat orientation, such as that between Arg554 and Asn546. There are a few intraprotomer interactions with loop2 that mimic those in the inclined orientation, especially from Tyr550 to Lys121 and Phe122 (Table S3, Fig. S11a; upper panel), but several others are not observed in the flat orientation, e.g., Ile549 to Phe122. Notably, there are no frequent contacts between the C-terminal tail and residues in loop4, unlike in the inclined orientation. Together these differences in intraprotomer contacts reflect that the start of the C-terminal helix is rotated outward toward the periphery of the trimer compared to a more ‘tucked in’ position in the inclined conformation (Fig. 1).

The flat orientation of the C-terminal helix allows many interprotomer contacts, but these differ substantially from those seen in the inclined orientation. Importantly, we do not observe contacts between any C-terminal segment and residues in the counterclockwise protomer, indicating a strong directionality of the interactions in the flat C-terminal orientation.

In the clockwise direction, the C-terminal helix interacts with different residues in the latter part of loop2 than in the inclined conformation. Notable interactions involve Ala133, Pro134, Glu135, Phe136 and Arg137, which interact with Ala564 and four different Arg residues (Table S3, Fig. 8c). Moreover, there are entirely new interactions with loop6; in particular, Arg576 forms frequent contacts with Ile298, Gly299 and Lys300 (Table S3, Fig. 8c, Fig. S11a; lower panel). Both loop2 and loop6 are directly connected to membrane-spanning helices (TM3 and TM7, respectively) that undergo major conformational changes upon closure of the cytoplasmic pathway, according to comparison between structures of different states in the transport cycle(10). Preserving these contacts in the flat orientation of the C-terminal helix is therefore likely to favor the observed inward-open conformation of the transport domain. Release of the C-terminal tail from those interactions may increase the probability of the inward-to outward-open conformational transition that is required during the betaine transport cycle. Consequently, these interactions are good candidates for hallmarks of a downregulated state.

**Figure 8.**
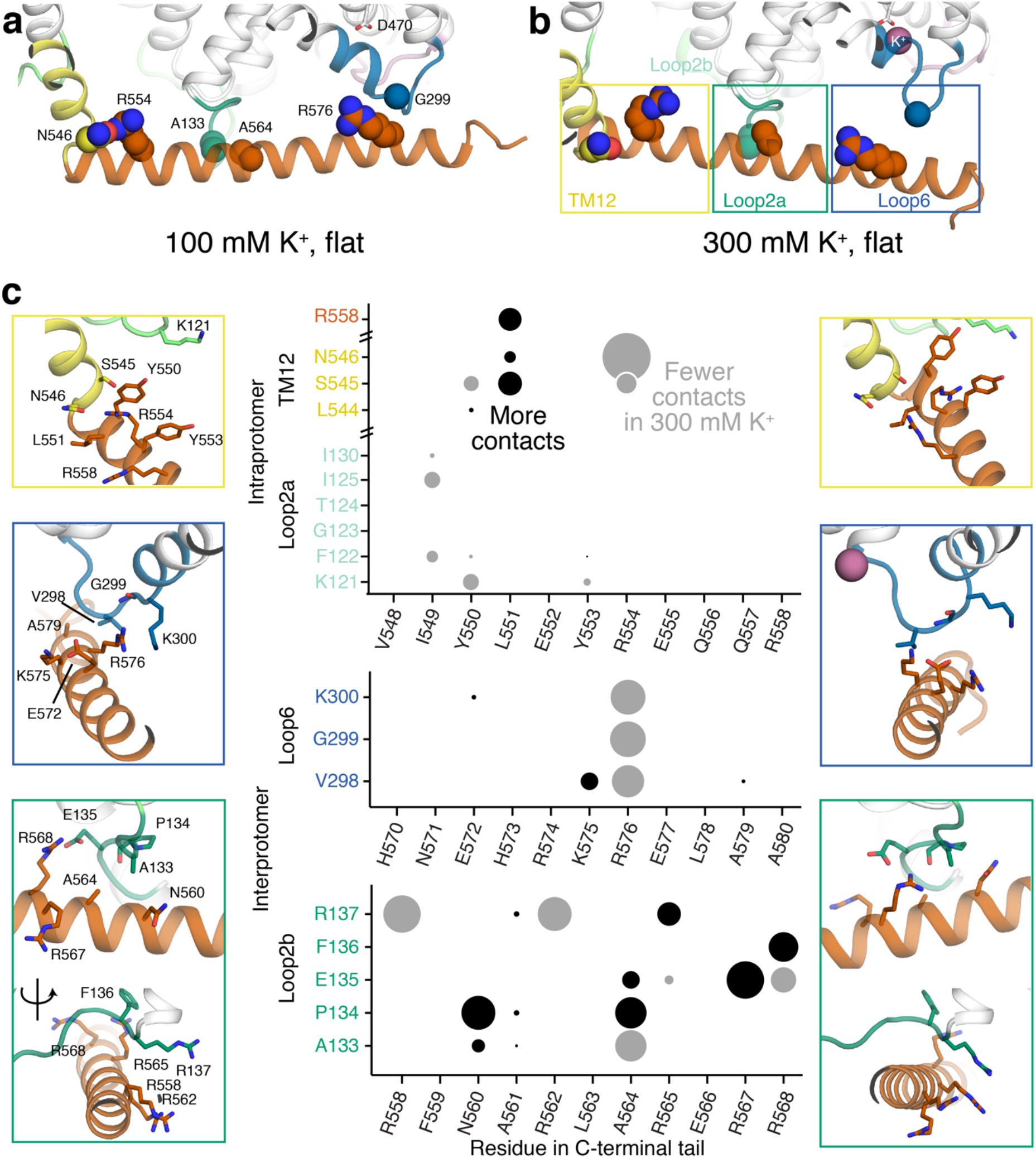
Contacts between the C-terminal tail of BetP and the rest of the trimer. **(a, b)** Contacts sampled during simulations of the flat orientation within either 100 mM (a) or 300 mM K^+^ (b). Interactions involve three distinct segments of the C-terminal tail. Boxes indicate the three regions in the close-ups in panel (c). The protein is shown as cartoon helices and colored according to Fig. 1. Ions and key interacting residues are shown as spheres. **(c)** In the close-up figures, pairs of residues forming contacts for >20% of the simulations are shown as sticks. The bubble plot illustrates each contact in these segments, and whether it is formed more frequently (black) or less frequently (gray) in the simulation with a higher concentration of K^+^. The size of each circle reflects the magnitude of the change.

#### The flat C-terminal conformation is stable in the presence of a high potassium concentration

Regulation of the transporter is thought to involve changes in the conformation of the C-terminal tail (38). We therefore examined whether K^+^ ions can disrupt, or otherwise influence, the contacts that keep the C-terminal helix flat against the BetP transporter domain, by simulating this conformation in the presence of 300 mM K^+^. However, throughout these five simulations, which amounted to 43.2 µs, considering each C-terminal helix as an independent entity, the C-terminal tail remained relatively stable according to the RMSD (Fig. 4b, Fig. S8f-j), the EM score (Fig. 5b, Fig. S10f-j), and the helicity (Fig. 4d, Fig. S9f-j). In addition, in the presence of a high concentration of K^+^, the helix remained parallel to the membrane (Fig. 6d). Similar to the simulations with 100 mM K^+^ (Fig. 7a, 7c) the C-terminal helix pivots somewhat on the surface of the trimer (Fig. 7b, 7d). We note that the distribution of pivot angles appears broader in the presence of 300 mM K^+^ (Fig. 7d), especially in the direction of the lipid bilayer; however, the most extreme values were only observed in individual simulation trajectories, limiting our ability to make robust conclusions about the effect of higher K^+^ concentration on tail orientation.

#### Potassium ions do not bind directly to the C-terminal segment, even at a high concentration

The fact that the global orientation of the C-terminal tail is similar in the simulations with both K^+^ concentrations suggests that K^+^ ions are not directly interfering with its ability to contact the cytoplasmic surface of the transport domain. Indeed, none of the cytoplasm-facing residues bound to K^+^ ions for a substantial proportion of the trajectories with the flat orientation and 100 mM K^+^ (Fig. S12). Nevertheless, K^+^ ions were present in the crevice adjacent to Asp470 in 7% of the trajectories with 300 mM K^+^, on average, leading to a distinct signal in the occupancy maps (Fig. 3d). Interestingly, the occupancy of this crevice with the flat C-terminal tail orientation is less than the ∼21% occupancy observed for the inclined orientation of the C-terminal tail at the same concentration (Fig. 3b). This discrepancy in occupancy between orientations cannot reflect direct K^+^ binding, since the C-terminal tail does not directly contact the ion or Asp470 in either orientation (Fig. S11, S12). However, as mentioned above, the C-terminal helix interacts frequently with loop6 in the flat orientation (Table S3, Fig. 3f, Fig. 8, Fig. S12) but not in the X-ray derived simulations (Table S3, Fig. 3e, Fig. S6).

One possibility is that the effect of this interaction between the C-terminal helix and loop6 propagates to the nearby crevice, reducing the affinity of Asp470 for cations and resulting in fewer binding events. Unfortunately, loop6 is extremely dynamic, hindering a robust comparison of its conformational ensembles in the two K^+^ concentrations. Nevertheless, it is reasonable to expect that binding of cations to Asp470 would result in fewer contacts between the cytoplasmic loops and the C-terminal tail. Indeed, compared with the simulations carried out with 100 mM K^+^, all three residues in loop6 make substantially fewer contacts with Arg576 at the higher K^+^ concentration (Fig. 8c).

Perhaps surprisingly, according to the helix orientation analysis described in the previous section, any changes in the contacts with loop6 do not appear to be associated with large movements of the helix. Nevertheless, there are other changes of around the same magnitude further upstream. For example, the mid-section of the C-terminal helix (residues 558-568), which forms many hydrophobic contacts with residues 133-137 in loop2 (Table S3), appears to shift its position by around 1-2 residues in the presence of 300 mM K^+^ (Fig. 8). For example, Arg558 and Arg562 lose contacts with Arg137, handing it off to Arg565. At the same time, Ala564 loses contacts with Ala133, but gains contacts with Pro134. Further upstream, close to the start of the C-terminal helix, there are additional changes. Most notably, Leu551 interacts more frequently with Ser545 in TM12, while Arg554 loses contacts with Asn546. All these small changes occurring simultaneously could provide a means by which K^+^ binding at the crevice near Asp470 might prime the transporter for activation events in which the C-terminal tail peels away from loop6 and loop2b, thereby releasing TM3 and TM7. At this point it is not clear whether, upon activation, the tail adopts a single preferred orientation like those in the X-ray structures, or if it samples many conformations involving interactions with lipids, the N-terminal tail, and/or loop2a.

To conclude, our simulations support the notion that the flat orientation represents a likely state under non-activating conditions, even in the absence of the N-terminal tail. Furthermore, this flat orientation is best described as a conformational ensemble encompassing various pivoting and rotating movements of the helix over the surface of the trimer and toward the lipid bilayer. Despite these dynamics, however, the C-terminal tail maintains many hydrophobic and charged contacts that together may be important for downregulation of the transport domain.

### Modeling of cytoplasmic tails on the full-length BetP trimer

Considering both the EM and simulation data, it seems clear that having the C-terminal helix flat against the protein periphery is a preferred arrangement for BetP under non-activating conditions. This finding provides the foundation for exploring the conformational space of the remaining missing ^piece^ of the BetP protein, namely the N-terminal tail. To determine the accessible conformational space of the N-terminal tail, we used *de-novo* modeling in the context of the EM-based BetP structure derived from our simulations. We used the FloppyTail algorithm (31) of the Rosetta package, which is designed to explore conformational space of flexible protein regions and not to predict a single “true” structure.

#### Remodeling the C-terminal helix with the de-novo protocol generates native-like structures

Before modeling the N-terminal tail, we tested the ability of this protocol to generate a BetP terminal tail by re-modeling the C-terminal helix and comparing against the EM structure. Specifically, we generated 100,000 *de-novo* models of the C-terminal tail in the context of the trimer.

Most of the C-terminal tail models obtained with FloppyTail are helical (Fig. S13b), indicating that this protocol can reliably create secondary structure, where needed. Even though most of these models adopt orientations that differ substantially from the EM structure (Fig. S13a), a few models have RMSD values <5 Å, thus reproducing the flat, extended C-terminal helix conformation. Unfortunately, given the limited degree of packing between the terminal tails and the rest of the protein, there is no suitable energy function for reliably identifying the correct orientations from among the other possibilities. Accordingly, when modeling either tail, we do not expect to be able to identify a single “true” conformation from those present in the sampled models. Instead, to understand whether this *de novo* approach can be used to gain molecular insights into BetP regulation we compared the predicted C-terminal tail contacts with those observed during our tens of microseconds of simulations.

As expected, the total count of contacts over the 100,000 models (Fig. S14) is lower than in the MD simulations (Fig. S11), since not all the C-terminal tail models are oriented horizontally on the surface of the trimer. Nevertheless, both in the models and in the simulations the C-terminal tail interacts consistently with loop2 of the same protomer, and of the adjacent protomer in the clockwise position (Fig. S11, S14). However, some contacts formed by the C-terminal tail in the simulations (especially with loop6 in the clockwise protomer, Fig. S11), are not found in any of the modeled C-terminal. *Vice versa*, a few contacts were found in the models - between the C-terminal tail and the C-terminal helix of the counterclockwise protomer – that were not observed in the simulations. This discrepancy is likely due to the larger (although, probably discontinuous) conformational space sampled in the models. In summary, with this *de-novo* strategy, we can reliably predict the secondary structure of a BetP terminal tail and the full set of models is likely to include native states. In addition, some, but not all, of the predicted contacts are likely to represent native interactions. Thus, the aggregate features of the N-terminal tail models are likely to be informative.

#### N-terminal models are diverse, but they share common secondary structure

After producing 100,000 models of the 55-residue long N-terminal tail, we clustered them based on structural similarity, to assess whether one conformation was preferentially sampled. However, the clustering analysis showed that the conformational space is very wide; there is no clear set of consistently sampled structures (Fig. 9a). Despite this conformational diversity, the secondary structure of the N-terminal tail models is similar across the whole set (Fig. 9b), with two well-defined α-helices covering residues 20-27 and 42-49. Allowing for other types of helices, these structured segments could be considered to encompass residues 5-31 and 33-50, or around 64% of the tail.

**Figure 9.**
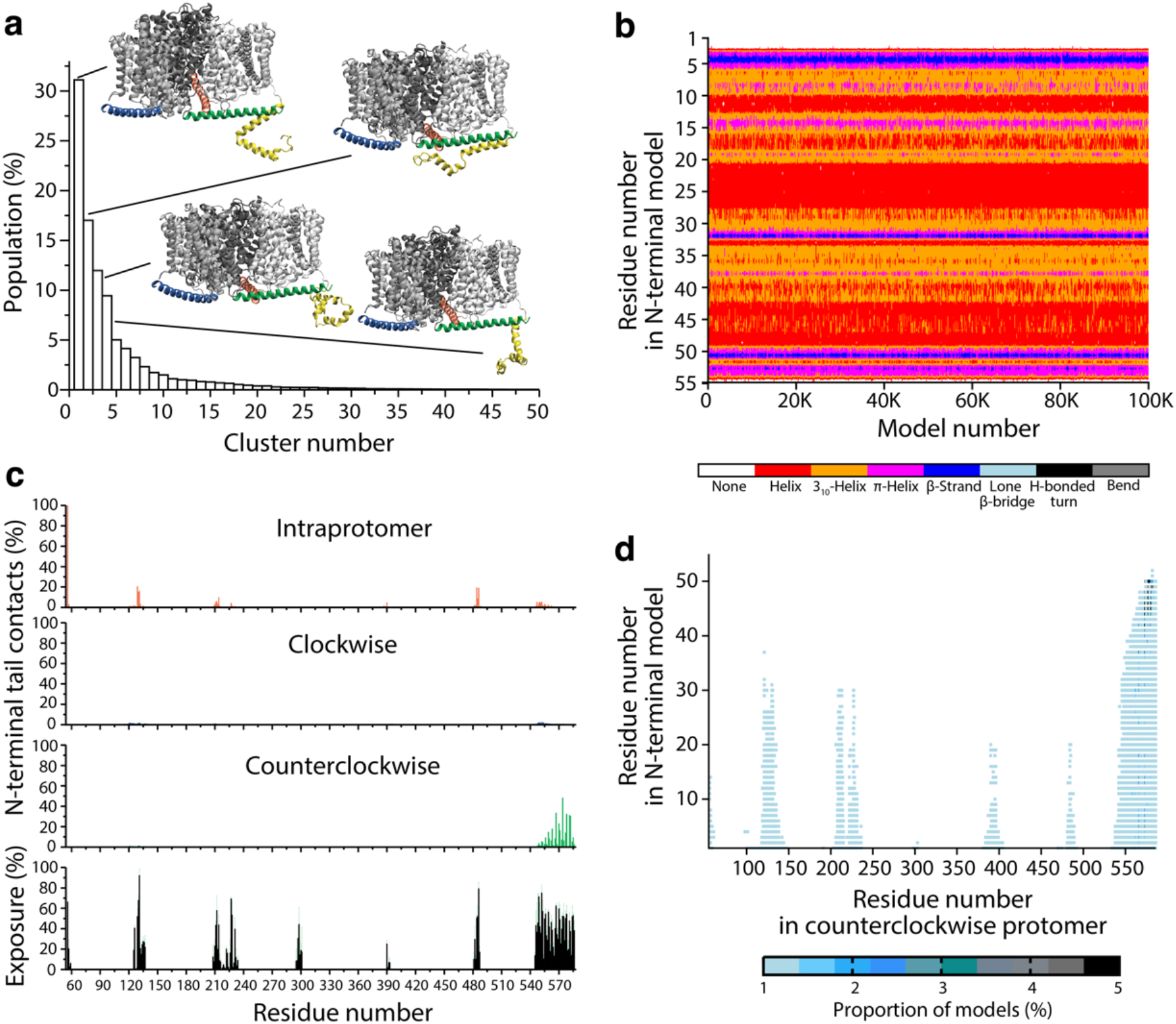
Structural features of the *de novo* predicted N-terminal tail of BetP. **(a)** Clusters of 100,000 models of residues 1 to 55 of the N-terminal tail, based on their Cα RMSD. For the four most-populated clusters, the structure at the center of that cluster is shown as cartoon helices, viewed from the plane of the membrane. The transmembrane segments are colored three shades of gray, the C-terminal segments are orange, blue and green, and the predicted N-terminal segment is colored yellow. **(b)** The secondary structure for all 100,000 N-terminal tail models, considering residues 1 to 55, computed using DSSP, plotted for every residue using the indicated color scheme. **(c)** Contacts of the modeled N-terminal tail residues 1 to 55 with individual residues in each of the three protomers of BetP, either intraprotomer (red bars), or interprotomer in the clockwise (blue bars), or counterclockwise (green bars) direction, assuming a view of the trimer from the cytoplasm. Contacts are defined as any two non-hydrogen atoms within 4.2 Å. Bottom panel: Available contact surface in the cytoplasmic regions of the BetP trimer, excluding the N-terminal segment. Data is plotted as the mean (black) and standard deviation (green error bars) over the three protomers of the solvent-accessible surface area, relative to the maximum possible for each amino acid type in a GXG tripeptide. **(d)** Contacts between any residue of the modeled N-terminal tail (residues 1 to 55) and residues in the counterclockwise protomer. Points are colored according to the number of models in which that contact was observed as indicated by the legend.

#### The N-terminal tail models do not interact with the adjacent clockwise protomer

The N-terminal tails are connected to TM1, which is located on the extreme periphery of the trimer and relatively far from the trimer interface (Fig. 1 and Fig. 8a-b), raising the question of whether the N-terminal segments ever interact with the adjacent protomers. We therefore measured the contacts formed by any N-terminal tail model with the rest of the trimer. Despite the presence of several solvent-accessible regions on the cytoplasmic surface of the trimer (Fig. 9c, bottom panel), only a few of those regions form contacts with the modeled N-terminal tails (Fig. 9c, upper panels). Some intraprotomer contacts were observed with residues in loop2 (around residue 130) and loop10 (around residue 480), but most contacts are formed between the N-terminal tail and the C-terminal tail of the adjacent protomer in the counterclockwise direction (Fig. 9c-d). Surprisingly, none of the models contain contacts between the N-terminal segments and the clockwise protomer, even though, as mentioned above, the *de novo* technique used here is likely to sample a conformational space wider than the true accessible space of the terminal tails.

#### Increasing the length of the modelled segment produced models with similar features

According to our benchmark in the C-terminal tail, the contact predictions may not be as reliable as the secondary structure of the models. We therefore wondered whether the lack of interaction of the N-terminal tail with the clockwise protomer reflects the choice of anchor point from which the N-terminal portion was modeled, i.e., residue 56, which was selected because it is present in most crystal structures of BetP. To test this possibility, we remodeled the N-terminal segment using the entire 60-residue segment up to the first helical residue of TM1. These models are also very diverse (Fig. S15a), but nevertheless exhibit similar secondary structure (Fig. S15b), with the same two helices as observed in the models generated for only the first 55 residues (Fig. 9b). Remarkably, also in this case, a negligible number of the 100,000 models form contacts with the clockwise protomer (Fig. S15c-d), supporting the notion that there is a directionality in the N-terminal tail interaction with the adjacent C-terminal helix.

#### Unstructured models of the N-terminal tail mainly contact the C-terminal helix of the counterclockwise protomer

The contacts adopted by the N-terminal tail models are governed by two factors: (i) the energetics of those interactions, or (ii) the steric and geometric constraints of the internal structure of the tail. In particular, the presence of helical segments shortens the available conformational space of the N-terminal tails. To further assess the robustness of the predicted interactions in the models, we built another set of 100,000 models of the N-terminal segment in which we did not use fragments derived from known crystal structures to guide the backbone conformational space. This approach reduced the formation of secondary structure elements. As expected, the sampled conformations are very diverse (Fig. S16a), with few and short secondary structure elements (Fig. S16b). However, even using this extreme approach, most of the N-terminal tail contacts involve the C-terminal tail in the counterclockwise protomer and almost none contact the clockwise protomer (Fig. S16c-d).

In summary, the results of these three sets of models and the EM-derived simulations, provide strong support for the possibility of crosstalk between the N- and C-terminal regulatory tails of BetP, and moreover, that this crosstalk has bi-directionality, i.e. involves the tails from both protomers on either side of a given subunit. As mentioned previously, the N-terminal tail is probably disordered to some extent, precluding its complete resolution in the EM structure. However, the EM data does reveal a distinct density connected to TM1 that points towards the counterclockwise C-terminal tail only (see Fig S9d in (12)), supporting the interaction model proposed by the de-novo modeling. In this model, each C-terminal helix contacts several different important regulatory segments, namely loop2 of its own protomer, as well as cytoplasmic loops (loop2 and loop6) and the N-terminal tail of the clockwise protomer (Fig. 8, 9, S15, S16).

#### Two major modes of interaction predicted for the N-terminal tails

The consistency of the predicted contacts across different modeling protocols suggests that some of the specific residues involved may be important for regulation. We therefore analyzed the predicted pairwise interactions between the N- and C-terminal tails. In all three sets of models, the most likely contacts involve residues 42-50 of the N-terminal tail, a Glu-rich region, which frequently forms salt-bridge interactions with the basic residues in the C-terminal helix (especially Arg574, Lys581 and Arg582; Fig. 9d, Fig.15d, Fig.16d). For example, a Glu50-Arg582 contact is observed in 16% of the 55-residue long N-terminal models. The predominant role of residues 42-50 is perhaps puzzling since deletions of the first 20 residues of the N-terminal tail showed the same regulatory effect as the truncation of the whole domain (7), implying that only residues 1-20 are essential for the N-terminal modulation of BetP activity. We therefore considered the possibility that the crosstalk between the cytoplasmic tails is not occurring via a few strongly-interacting pairs of residues but is instead an accumulation of interactions along the two tails. As previously mentioned, the models of the N-terminal tail consistently predicted two helical regions, separated by an unstructured, potentially hinge-like region. Thus, these helices might act as two independent entities that either contact each other, if folded together, or contact the C-terminal tail, if extended. To test this hypothesis, we plotted the contacts formed between the N- and C-terminal tails against the interactions between the two N-terminal helices for each model (Fig. 10a). Most of the N-terminal tail models either form contacts with the C-terminal helix or contain contacts between the two N-terminal helices, or neither of these two options. Only rarely can the N-terminal helices contact both the C-terminal tail and each other.

**Figure 10.**
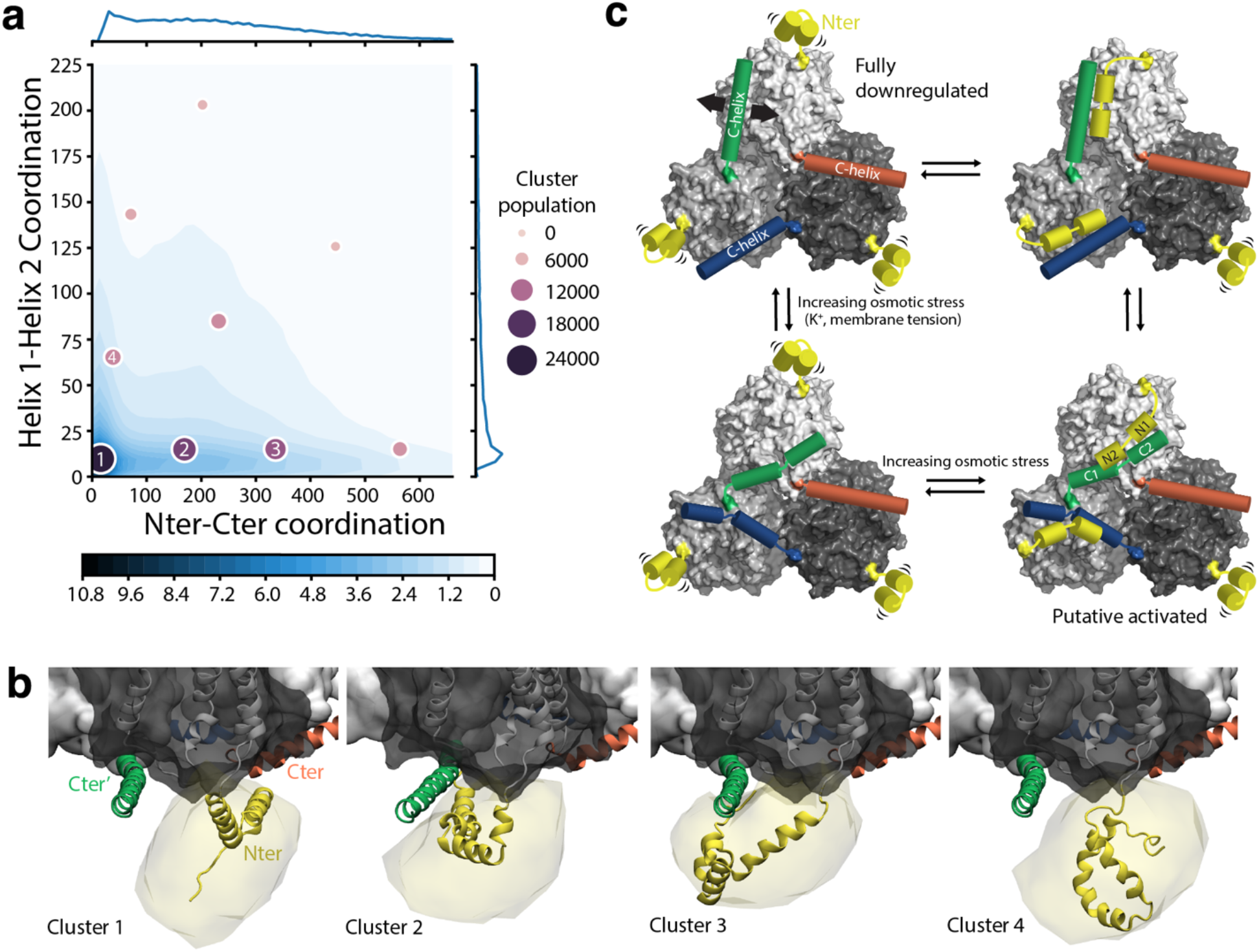
Interactions of the modeled N-terminal tail and a hypothesis for their role in activation. **(a)** Contacts within the N-terminal tail (residues 1-55) and between the N-terminal tail and the C-terminal tail (residues 548-586) plotted as the proportion of the 100,000 de novo models (blue scale-bar). Contacts are defined using a coordination number (see Methods) for non-hydrogen atoms of the C- and N-terminal tails versus a coordination number between N-terminal helix 1 (residues 10-31) and helix 2 (residues 36-49). Clusters based on these two variables are mapped onto the 2D plot, with the size of each cluster indicated by the size and purple shading of the circles. **(b)** Structures of the four most populated clusters in the space of these two variables, viewed along the plane of the membrane. The variance of the models is indicated using surface representation (yellow), while the N-terminal tail model with the greatest degree of helicity and with the corresponding coordination numbers, is shown as yellow cartoon helices. The transmembrane segment of the protomer containing the modelled N-terminal tail is shown as a dark gray transparent surface, with helices in white (transmembrane segments) and orange (C-terminal tail). The other two protomers are shown as white surfaces, with the C-terminal tail of the counterclockwise protomer shown as a green cartoon. **(c)** Schematic of likely intra- and inter-tail interactions predicted by the modeling. The fully downregulated state (top left) is expected to resemble that observed in EM measurements in amphipols and low K^+^ concentrations, which is stable in MD simulations. The lack of resolution of the N-terminal tail under these conditions suggests that it is either fully unfolded or connected by a flexible linker, as in clusters 1 and 4. Activation may involve a shift in the conformation of one or more C-terminal helices away from loop6 and loop2b, perhaps toward the trimer midpoint, similar to its orientation in X-ray crystal structures; in this conformation, the helical segment beyond the breakage point (C2) is likely to be more dynamic (bottom panels). Additionally, an extended conformation of the N-terminal helix is predicted to form contacts with the C-terminal domain, as in clusters 2 and 3, suggesting activated state configurations such as those on the right.

To obtain a molecular-level picture of this effect, we clustered the models in the space of these two contact variables. The most populated cluster (cluster 1 in Fig. 10a-b) represents extended N-terminal tails occupying spaces away from the rest of the BetP trimer. We consider this an unlikely possibility since truncations alter regulation profiles(5). In the second and third most populated clusters, the structures lack contacts between the N-terminal helices but have many interactions with the C-terminal tails; here, the N-terminal tails are more extended and lie closer to the C-terminal tail. Lastly, the N-terminal models of the fourth most-populated cluster are more compact, as if forming a separate domain involving intra-tail contacts and lacking interactions with the C-terminal tail.

## Discussion and Conclusions

In this work we set out to gain understanding of the regulatory mechanism of a secondary active transporter by determining the structural features of the N- and C-terminal domains of BetP, combining available structural data with µs-scale simulations and *de novo* modeling.

Of the two domains, more structural information is available for the C-terminal segment, which is also essential for activity regulation. Hence, we first used molecular dynamics simulations to assess the stability of two observed orientations of this segment: one derived from X-ray crystallographic data and another from EM data. The C-terminal domain derived from the EM data obtained in low K^+^ concentrations is stable for up to 8 µs in a PG lipid bilayer, suggesting that it corresponds to a down-regulated conformation of the transporter. By contrast, the simulations of the conformation observed in the X-ray structure gave a more mixed picture: the most C-terminal portion of this helix, which is involved in contacts with other trimers in the crystal lattice, was not stable in any of our simulations. No unique conformation, nor even a distinct conformational ensemble, was obtained for this segment. Rather, irreversible conformational drift away from the initial crystal structure orientation was observed, resulting in very different configurations for each simulation. These configurations appear to be stabilized by protein-protein interactions that seemingly trap the C-terminal segments and prevent them from sampling a true energy minimum. Notably, however, the portion of the C-terminal segment directly linked to the transmembrane region partially retains the conformation observed in the crystal structure. In this orientation, this segment spans the interior cytoplasmic surface of the trimer far from the lipid bilayer and has been suggested to reflect an activated conformation, because (a) the trimer arrangement is asymmetric, and (b) contains contacts involving residues known to be essential for regulation (12).

We attempted to explore the possible active state of BetP in our simulations by increasing the concentration of K^+^ ions, a known regulatory stimulus, to 300 mM. On the microsecond timescale used in our simulations, only local conformational changes of the C-terminal helix conformation were observed, whereas a transition to an activated state might be expected to involve a more major conformational change, for example toward an inclined conformation. This lack of sampling of other states may reflect insufficient simulation time, but might also reflect that in the absence of the N-terminal domain BetP may require a higher concentration of K^+^ for activation than the full-length protein (5, 7).

It is worth noting that PG lipids are associated with a shift toward higher values of the osmotic stress profile, potentially strongly downregulating the transporter compared to *E. coli* cells or a *C. glutamicum* lipid mixture (39). Our choice of lipid was driven by the fact that PG lipids associate tightly with BetP even during purification and detergent solubilization(14, 40). Moreover, the use of a single lipid component avoids artefacts in the protein conformational ensemble due to limited convergence of the lipids in a mixed bilayer during µs-scale simulations. However, future studies should investigate the behavior of complex mixtures on BetP given their key role in regulation.

Unexpectedly, the simulations revealed repeated binding of K^+^ ions to the transporter domain, in a small pocket adjacent to the cytoplasmic pathway comprising Asp470 in TM10 on one side and loop6 (between TM6 and TM7) on the other side. Loop6 is notable for the numerous contacts it forms with the C-terminal helix in the downregulated conformation of the protein (Table S3, Fig. 8). Asp470 was previously predicted to experience a shift in protonation state depending on the conformation of the transport domain, and on the presence of a Na^+^ ion bound at the so-called Na2 site in the core of the transporter (16). In the inward-facing conformation used here, Asp470 was predicted to be deprotonated at pH 7 and therefore, in our simulations of the trimer, Asp470 was *a priori* assigned to be deprotonated, based on which one might expect that cations will inevitably bind there at the high concentrations of K^+^ used (300 mM). Nevertheless, no other negatively-charged residues on the surface of the protein experienced a similar frequency of cation interactions. We note that the cation interactions with Asp470 were not specific to K^+^ ions, as Na^+^ ions were also found to bind in this pocket, albeit with a lower rate, perhaps consistent with its lower concentration (200 mM). Similarly, the aforementioned shifts in predicted protonation states indicate that protons might also compete for this site. Indeed, at pH 5.5, which is required for crystallization, we expect a significant population of the protonated form of Asp470, which may explain why no density for Rb^+^ was observed at this region in crystals obtained with 300 mM RbCl (12). Taken together with the high basal levels of K^+^, low levels of Na^+^, and a neutral pH in the cytoplasm of *C. glutamicum* (41, 42), we suggest that Asp470 could play a key role in activation by K^+^ ions.

Given the location of this pocket immediately adjacent to the highly basic C-terminal helix (Fig. 3f, 8), we speculated that cation binding reduces the likelihood of the “flat” orientation of the C-terminal helix and therefore leads to activation by releasing the contacts with loop2b immediately prior to TM3, and thereby enabling closure of the cytoplasmic pathway. However, we were unable to establish systematic effects of K^+^ binding on the global conformation of the C-terminal tail or of loop6. This is perhaps because the C-terminal tail(9) would be required to release numerous (albeit generally weak) interactions with residues in loop2 and loop6 to escape the downregulated conformation, likely a slow process relative to the microseconds of sampling achieved here, especially given the limited occupancy of the putative K^+^ site. Therefore, it remains to be established, on the molecular level, whether and how direct K^+^ binding at this site provides the foundation for accelerated turnover by the transport domain.

Building on our understanding of the C-terminal helix orientation in the downregulated state we sampled the possible conformations of the N-terminal tail, which has a modulatory action on the response of BetP to osmotic stress. The absence of structural data for the N-terminal tail, even in the EM experiment where it is present in the protein construct, could suggest that it forms no consistent structure nor contacts with the rest of the protein. However, previous studies revealed that modified BetP proteins with truncations of the first 20, 29, 32, 49, 52, 53 or 60 residues (the latter of which removes the entire N-terminal segment) all require higher osmolalities for activation, and have a reduced maximum velocity (7, 9), indicating either that the downregulated state(s) is further stabilized by the absence of this domain, or that the activated state(s) are more readily formed in its presence. Moreover, peptides with the N-terminal sequence compete with the native attached N-terminal domain to limit the ability of the transporter to reach maximal turnover levels. The nature of its role, from a structural perspective, is therefore puzzling. We can rule out that N-terminal segment interacts primarily with the membrane surface, based on surface plasmon resonance analysis (9), as well as high density of negative charges in both the lipid headgroups and the N-terminal tail. Therefore, we think it likely that the N-terminal tail does interact with the rest of the protein at some stage during activation, and presumably with the C-terminal tail(s), given its density of positive charges and observed interactions in peptide arrays (9). Here, instead of proposing a unique model of the structure of the N-terminal domain, we identified common features across thousands of different models, repeatedly challenging our findings by varying the modeling strategy. The models consistently indicated interactions between the N-terminal tail of one protomer with the C-terminal helix of the adjacent protomer in the counterclockwise position (Fig. 9c), a notion supported by the low resolved density assigned to the first portion of the N-terminal tail in the EM structure. By contrast, the most stable configuration of the C-terminal tail, the one derived from the EM data, interacts with the adjacent protomer in the clockwise direction. Therefore, each protomer contacts both of its neighbors in the trimer, one through its N-terminal domain and the other via the C-terminal helix, indicating a strong bi-directional interplay between the three subunits (Fig. 10c).

That truncations of the first 20, 32, 49 or 60 residues all led to similar shifts in activation curves relative to the WT protein (7) might be interpreted as indicating that only the first 20 residues form important contacts with the rest of the protein. Instead, we suggest here, based on the N-terminal tail models, that the complex between the N- and C-terminal domains involves numerous, weak contacts, including those in the first 20 residues. Consequently, elimination of a few residues in the beginning of the N-terminal segment could disrupt the interaction between the N- and C-terminal domains. A need for many, weak, cooperative residue-residue interactions between the terminal domains may be an advantage for the regulation mechanism. In this way, disrupting only a few residue-residue contacts (e.g., by competition with cations or changes in local membrane structure) may suffice to completely disengage the N- and C-terminal complex, which in turn may activate BetP transport (Fig. 10).

Taken together, although additional mechanistic details remain to be established, especially with respect to the role of the membrane in osmoregulation, these computational studies enhance our understanding of the structure and dynamics of the BetP trimer and of its regulatory domains.

A similar modeling strategy to that used for the N-terminal domain was used previously for the cytoplasmic N- and C-terminal tails of the human serotonin transporter (hSERT) to predict that they contain conserved secondary structural elements (43). Here, using FloppyTail, we could explicitly consider the possible unstructured nature of the N-terminal segment and its context in the trimer, allowing us to predict interaction patterns between the modeled segment and the rest of the protein. Such interactions between cytoplasmic segments and the transmembrane core are likely to be of interest for many other membrane proteins. In particular, terminal segments are often not resolved in structural studies, but can play important roles in membrane trafficking, activity regulation, posttranslational modification, etc (44-47), that may involve interactions with the rest of protein or with the lipid bilayer. Therefore, although the proposed regulatory interactions presented here is likely to be specific to BetP, we expect that similar computational strategies will be useful for obtaining structural insights into the regulatory terminal segments of other membrane proteins.

## Acknowledgements

This research was supported by the Division of Intramural Research of the NIH, National Institute of Neurological Disorders and Stroke, NS003139 and the Deutsche Forschungsgemeinschaft (DFG, German Research Foundation) through SFBs 699 and 1350. We thank Drs. José Faraldo-Gómez, Edward Lyman and Richard Venable for useful discussions and Dr. Richard Venable for sharing scripts and analysis of their POPG simulations. We used the computational resources of the National Institutes of Health High Performing Computing cluster, Biowulf. Anton2 computer time was provided by the Pittsburgh Supercomputing Center (PSC) through Grant R01GM116961 from the National Institutes of Health. The Anton2 machine at PSC was generously made available by D.E. Shaw Research.

## References

1. Csonka LN, Hanson AD. Prokaryotic osmoregulation: genetics and physiology. Annu Rev Microbiol. 1991;45:569–606.

2. Kramer R, Morbach S. BetP of Corynebacterium glutamicum, a transporter with three different functions: betaine transport, osmosensing, and osmoregulation. BBA-Bioenergetics. 2004;1658(1-2):31–6.

3. Ziegler C, Morbach S, Schiller D, Kramer R, Tziatzios C, Schubert D, et al. Projection structure and oligomeric state of the osmoregulated sodium/glycine betaine symporter BetP of Corynebacterium glutamicum. J Mol Biol. 2004;337(5):1137–47.

4. Tsai CJ, Ziegler C. Structure determination of secondary transport proteins by electron crystallography: Two-dimensional crystallization of the betaine uptake system BetP. J Mol Microb Biotech. 2005;10(2-4):197–207.

5. Ressl S, van Scheltinga Act, Vonrhein C, Ott V, Ziegler C. Molecular basis of transport and regulation in the Na+/betaine symporter BetP. Nature. 2009;458(7234):47–U1.

6. Perez C, Khafizov K, Forrest LR, Kramer R, Ziegler C. The role of trimerization in the osmoregulated betaine transporter BetP. EMBO Rep. 2011;12(8):804–10.

7. Peter H, Burkovski A, Kramer R. Osmo-sensing by N-and C-terminal extensions of the glycine betaine uptake system BetP of Corynebacterium glutamicum. J Biol Chem. 1998;273(5):2567–74.

8. Rubenhagen R, Morbach S, Kramer R. The osmoreactive betaine carrier BetP from Corynebacterium glutamicum is a sensor for cytoplasmic K+. Embo J. 2001;20(19):5412–20.

9. Ott V, Koch J, Spate K, Morbach S, Kramer R. Regulatory Properties and Interaction of the C-and N-Terminal Domains of BetP, an Osmoregulated Betaine Transporter from Corynebacterium glutamicum. Biochemistry. 2008;47(46):12208–18.

10. Perez C, Koshy C, Yildiz O, Ziegler C. Alternating-access mechanism in conformationally asymmetric trimers of the betaine transporter BetP. Nature. 2012;490(7418):126–30.

11. Tsai CJ, Khafizov K, Hakulinen J, Forrest LR, Forrest LR, Kramer R, et al. Structural asymmetry in a trimeric Na+/betaine symporter, BetP, from Corynebacterium glutamicum. J Mol Biol. 2011;407(3):368–81.

12. Heinz V, Gueler G, Leone V, Madej MG, Maksimov S, Gaertner RM, et al. Osmotic stress response in BetP: How lipids and K+ team up to overcome downregulation. bioRxiv. 2022:2022.06.02.493408.

13. Waclawska I, Ziegler C. Regulatory role of charged clusters in the N-terminal domain of BetP from Corynebacterium glutamicum. Biol Chem. 2015;396(9-10):1117–26.

14. Koshy C, Schweikhard ES, Gartner RM, Perez C, Yildiz O, Ziegler C. Structural evidence for functional lipid interactions in the betaine transporter BetP. Embo J. 2013;32(23):3096–105.

15. Williams CJ, Headd JJ, Moriarty NW, Prisant MG, Videau LL, Deis LN, et al. MolProbity: More and better reference data for improved all-atom structure validation. Protein Sci. 2018;27(1):293–315.

16. Leone V, Waclawska I, Kossmann K, Koshy C, Sharma M, Prisner TF, et al. Interpretation of spectroscopic data using molecular simulations for the secondary active transporter BetP. J Gen Physiol. 2019;151(3):381–94.

17. Zhang L, Hermans J. Hydrophilicity of cavities in proteins. Proteins. 1996;24(4):433–8.

18. Staritzbichler R, Anselmi C, Forrest LR, Faraldo-Gomez JD. GRIFFIN: A Versatile Methodology for Optimization of Protein-Lipid Interfaces for Membrane Protein Simulations. J Chem Theory Comput. 2011;7(4):1167–76.

19. Trabuco LG, Villa E, Mitra K, Frank J, Schulten K. Flexible fitting of atomic structures into electron microscopy maps using molecular dynamics. Structure. 2008;16(5):673–83.

20. DiMaio F, Tyka MD, Baker ML, Chiu W, Baker D. Refinement of Protein Structures into Low-Resolution Density Maps Using Rosetta. J Mol Biol. 2009;392(1):181–90.

21. Phillips JC, Braun R, Wang W, Gumbart J, Tajkhorshid E, Villa E, et al. Scalable molecular dynamics with NAMD. J Comput Chem. 2005;26(16):1781–802.

22. Best RB, Zhu X, Shim J, Lopes PEM, Mittal J, Feig M, et al. Optimization of the Additive CHARMM All-Atom Protein Force Field Targeting Improved Sampling of the Backbone phi, psi and Side-Chain chi(1) and chi(2) Dihedral Angles. J Chem Theory Comput. 2012;8(9):3257–73.

23. Klauda JB, Venable RM, Freites JA, O’Connor JW, Tobias DJ, Mondragon-Ramirez C, et al. Update of the CHARMM All-Atom Additive Force Field for Lipids: Validation on Six Lipid Types. J Phys Chem B. 2010;114(23):7830–43.

24. Venable RM, Brown FLH, Pastor RW. Mechanical properties of lipid bilayers from molecular dynamics simulation. Chem Phys Lipids. 2015;192:60–74.

25. Jorgensen WL, Chandrasekhar J, Madura JD, Impey RW, Klein ML. Comparison of Simple Potential Functions for Simulating Liquid Water. J Chem Phys. 1983;79(2):926–35.

26. Darden T Y D L P. Particle mesh Ewald: An N.log(N) method for Ewald sums in large systems. J Chem Phys. 1993;98(12):10089–92.

27. Lippert RA, Predescu C, Ierardi DJ, Mackenzie KM, Eastwood MP, Dror RO, et al. Accurate and efficient integration for molecular dynamics simulations at constant temperature and pressure. J Chem Phys. 2013;139(16).

28. Predescu C, Lerer AK, Lippert RA, Towles B, Grossman JP, Dirks RM, et al. The u-series: A separable decomposition for electrostatics computation with improved accuracy. J Chem Phy 2020;152(8).

29. Kabsch W, Sander C. Dictionary of Protein Secondary Structure - Pattern-Recognition of Hydrogen-Bonded and Geometrical Features. Biopolymers. 1983;22(12):2577–637.

30. McGibbon RT, Beauchamp KA, Harrigan MP, Klein C, Swails JM, Hernandez CX, et al. MDTraj: A Modern Open Library for the Analysis of Molecular Dynamics Trajectories. Biophys J. 2015;109(8):1528–32.

31. Kleiger G, Saha A, Lewis S, Kuhlman B, Deshaies RJ. Rapid E2-E3 Assembly and Disassembly Enable Processive Ubiquitylation of Cullin-RING Ubiquitin Ligase Substrates. Cell. 2009;139(5):957–68.

32. Jumper J, Evans R, Pritzel A, Green T, Figurnov M, Ronneberger O, et al. Highly accurate protein structure prediction with AlphaFold. Nature. 2021;596(7873):583.

33. Li SC, Ng YK. Calibur: a tool for clustering large numbers of protein decoys. BMC Bioinformatics. 2010;11.

34. Fiorin G, Klein ML, Hénin J. Using collective variables to drive molecular dynamics simulations. Molecular Physics. 2013;111(22-23):3345–62.

35. Giorgino T, Laio A, Rodriguez A. METAGUI 3: A graphical user interface for choosing the collective variables in molecular dynamics simulations. Comput Phys Commun. 2017;217:204–9.

36. Biarnes X, Pietrucci F, Marinelli F, Laio A. METAGUI. A VMD interface for analyzing metadynamics and molecular dynamics simulations. Comput Phys Commun. 2012;183(1):203–11.

37. Becker M, Maximov S, Becker M, Meyer U, Wittmann A, Kramer R. Analysis of putative protomer crosstalk in the trimeric transporter BetP: The heterotrimer approach. Biochim Biophys Acta. 2014;1837(6):888–98.

38. Ge L, Perez C, Waclawska I, Ziegler C, Muller DJ. Locating an extracellular K+-dependent interaction site that modulates betaine-binding of the Na+-coupled betaine symporter BetP. Proc Natl Acad Sci U S A. 2011;108(43):E890–8.

39. Schiller D, Ott V, Kramer R, Morbach S. Influence of membrane composition on osmosensing by the betaine carrier BetP from Corynebacterium glutamicum. J Biol Chem. 2006;281(12):7737–46.

40. Tsai CJ, Ejsing CS, Shevchenko A, Ziegler C. The role of lipids and salts in two-dimensional crystallization of the glycine-betaine transporter BetP from Corynebacterium glutamicum. J Struct Biol. 2007;160(3):275–86.

41. Follmann M, Becker M, Ochrombel I, Ott V, Kramer R, Marin K. Potassium transport in corynebacterium glutamicum is facilitated by the putative channel protein CglK, which is essential for pH homeostasis and growth at acidic pH. J Bacteriol. 2009;191(9):2944–52.

42. Kramer R, Lambert C, Hoischen C, Ebbighausen H. Uptake of glutamate in Corynebacterium glutamicum. 1. Kinetic properties and regulation by internal pH and potassium. Eur J Biochem. 1990;194(3):929–35.

43. Fenollar-Ferrer C, Stockner T, Schwarz TC, Pal A, Gotovina J, Hofmaier T, et al. Structure and Regulatory Interactions of the Cytoplasmic Terminal Domains of Serotonin Transporter. Biochemistry. 2014;53(33):5444–60.

44. Foss SM, Li HY, Santos MS, Edwards RH, Voglmaier SM. Multiple Dileucine-like Motifs Direct VGLUT1 Trafficking. J Neurosci. 2013;33(26):10647–60.

45. Sweeney CG, Tremblay BP, Stockner T, Sitte HH, Melikian HE. Dopamine Transporter Amino and Carboxyl Termini Synergistically Contribute to Substrate and Inhibitor Affinities. J Biol Chem. 2017;292(4):1302–9.

46. Bianchi F, van’t Klooster JS, Ruiz SJ, Poolman B. Regulation of Amino Acid Transport in Saccharomyces cerevisiae.Microbiol Mol Biol R. 2019;83(4).

47. Mikros E, Diallinas G. Tales of tails in transporters. Open Biol. 2019;9(6).

