## Supplementary Materials for "Autoregulation of a trimeric transporter involves the cytoplasmic domains of both adjacent subunits"

### Supplementary Text

#### *Test of lipid dynamics on the Anton2 high performance supercomputer*

Since the simulation parameters of Anton2, which uses a unique, integrated molecular dynamics software engine, differ from those used to parametrize the Charmm36 force field [1], we assessed whether simulations of a pure POPG lipid bilayer carried out on Anton2 exhibited similar dynamics and ensemble properties to those carried out using the Charmm molecular dynamics package [1]. Our system reflected that used in the simulations of BetP, and contained 648 POPG molecules, with a water:lipid ratio of 45:1, 738 sodium ions and 90 chloride ions, giving a total of 170,604 atoms. The MD trajectory obtained using Charmm [2] was generously provided by Drs. Richard Venable and Richard Pastor.

We performed three different simulations using Anton2 and varied either the cutoff distance at which the van der Waals (vdW) potential was truncated (either 9 or 12 Å), or the treatment of long-range electrostatic interactions (either Gaussian Split Ewald (GSE) [3] or U-series [4]). All other parameters were selected automatically by the Anton2 "chooser" routine. In all simulations we used a 2 fs-long time step, and the MD integrator was accelerated using a reference system propagator algorithm (RESPA) to compute long-range forces every three steps. The temperature was set to 303 K, pressure was maintained at 1.0 bar with semi-isotropic coupling, and each simulation was 480 ns long. For reference, the Charmm simulations included 303 POPG lipids at a 45:1 water:lipid ratio, vdW forces were switched off smoothly at a distance between 8 and 12 Å, the particle-mesh Ewald method was used for long-range electrostatic interactions, and the integration timestep was 1 fs.

We first compared the average area per lipid in each simulation, which is 67.5 Å$^2$ in the reference Charmm simulation [2]. When using a similar 12 Å cutoff for the vdW interactions, the area per lipid in the Anton2 simulations differ by ~5 Å$^2$ (62.4 ± 0.6 Å$^2$ over the last 100 ns, **Fig. S2a**). However, for both Anton2 simulations carried out using the shorter (9 Å) cutoff, the area per lipid differs by only ~2 Å (65.8 ± 0.7 and 65.6 ± 0.6 Å$^2$), independent of the long-range electrostatics scheme used.

We also compared the deuterium order parameters for the POPG tails in the different simulations, defined as:

$$S_{CD} = \left| \frac{< 3cos^2\theta - 1 >}{2} \right|$$

where $\theta$ is the angle of the C–H vector of a given carbon with respect to the membrane normal. Again, using this metric the simulations generated with a 9 Å vdW cutoff (with either GSE or U-series electrostatics) most closely reproduce the results obtained with the Charmm program [2] (**Fig S2b-c**). Hence, all the simulations presented in this work were performed with a hard vdW cutoff of 9 Å and using U-series long-range electrostatics.

**Table S1.** Summary of molecular dynamics simulations of BetP, in microseconds

| **C-helix orientation** | **100 mM $K^+$** | **300 mM $K^+$** | **Total** |
|---|---|---|---|
| Inclined (X-ray) | 4.8, 2.4, 2.4, 2.4 | 4.8, 2.4, 2.4, 2.4 | 24.0 |
| Flat (EM) | 7.4, 2.4, 2.4, 2.4, 2.4 | 4.8, 2.4, 2.4, 2.4,2.4 | 31.4 |
| Both | 29.0 | 26.4 | 55.4 |

**Table S2.** Contacts between C-terminal tail residues and the rest of the trimer in the reference crystal structure of BetP

| | | Residues 548-564 of protomer A | Residues 548-564 of protomer B | Residues 548-564 of protomer C |
|---|---|---|---|---|
| Residues in the rest of the trimer | A | 121, 122, 125, 210, 544, 545, 546, 547 | | 568 |
| | B | | 122, 124, 125, 210, 544, 545, 546, 547 | 131 |
| | C | 131 | | 121, 122, 125, 210, 544, 545, 546, 547 |

Contacts are defined as non-hydrogen atoms ≤ 4.2Å apart in the crystal structure (PDB code 4C7R) and are separated according to whether the residues contacted by the C-terminal residues are intraprotomer (diagonals); or in a neighboring protomer in either the clockwise (gray) or counterclockwise (dark gray) position, as viewed from the cytoplasm.

**Table S3**. Contacts between C-terminal tail residues and the rest of the trimer in MD simulations

| C-tail | | Rest of protein | | Inclined 100 mM | Inclined 300 mM | Flat 100 mM | Flat 300 mM |
|---|---|---|---|---|---|---|---|
| Intraprotomer | | | | | | | |
| Val | 548 | Arg | 210 | 38.3 | 29.7 | 5.1 | 20.1 |
| Ile | 549 | Phe | 122 | 19.7 | 35.7 | 1.5 | 7.1 |
| Ile | 549 | Ile | 125 | 55.5 | 56.8 | 4.9 | 12.8 |
| Ile | 549 | Ile | 130 | 25.2 | 22.1 | 0.5 | 1.9 |
| Ile | 549 | Tyr | 206 | 27.0 | 23.4 | 1.1 | 7.4 |
| Ile | 549 | Arg | 210 | 83.0 | 69.9 | 4.3 | 12.0 |
| Tyr | 550 | Lys | 121 | 44.9 | 30.3 | 33.1 | 41.0 |
| Tyr | 550 | Phe | 122 | 87.6 | 73.3 | 48.5 | 49.3 |
| Tyr | 550 | Leu | 544 | 55.5 | 77.2 | 39.7 | 41.1 |
| Tyr | 550 | Ser | 545 | 58.0 | 71.1 | 58.3 | 50.6 |
| Leu | 551 | Ser | 545 | 26.4 | 18.9 | 47.7 | 60.0 |
| Leu | 551 | Asn | 546 | 30.3 | 22.2 | 64.6 | 69.9 |
| Leu | 551 | Arg | 558 | 6.7 | 7.3 | 20.3 | 32.2 |
| Tyr | 553 | Lys | 121 | 75.3 | 57.9 | 2.1 | 5.0 |
| Tyr | 553 | Phe | 122 | 10.1 | 40.1 | 0.0 | 0.2 |
| Arg | 554 | Ser | 545 | 4.1 | 1.7 | 48.4 | 36.0 |
| Arg | 554 | Asn | 546 | 0.2 | 0.0 | 32.2 | 6.0 |
| Interprotomer clockwise | | | | | | | |
| Gln | 557 | Thr | 124 | 6.2 | 27.3 | 0.0 | 0.0 |
| Arg | 558 | Asp | 131 | 27.8 | 28.4 | 3.5 | 7.7 |
| Arg | 558 | Arg | 137 | 0.0 | 0.0 | 27.0 | 6.6 |
| Asn | 560 | Ala | 133 | 0.0 | 0.0 | 69.0 | 75.6 |
| Asn | 560 | Pro | 134 | 0.0 | 0.0 | 11.1 | 29.6 |
| Asn | 560 | Tyr | 553 | 5.2 | 27.3 | 0.0 | 0.0 |
| Ala | 561 | Ala | 133 | 0.0 | 0.0 | 31.1 | 31.6 |
| Ala | 561 | Pro | 134 | 0.0 | 0.0 | 55.3 | 57.1 |
| Ala | 561 | Arg | 137 | 0.0 | 0.0 | 41.6 | 43.4 |
| Arg | 562 | Asp | 131 | 54.8 | 41.9 | 0.0 | 0.0 |
| Arg | 562 | Arg | 137 | 0.0 | 0.0 | 39.0 | 20.5 |
| Leu | 563 | Gln | 556 | 10.4 | 28.8 | 0.0 | 0.0 |
| Ala | 564 | Ala | 133 | 0.0 | 0.0 | 56.1 | 39.3 |
| Ala | 564 | Pro | 134 | 0.0 | 0.0 | 51.5 | 68.6 |
| Ala | 564 | Glu | 135 | 0.0 | 0.0 | 48.8 | 57.8 |
| Ala | 564 | Ile | 549 | 13.9 | 32.7 | 0.0 | 0.0 |
| Ala | 564 | Gln | 556 | 33.2 | 22.6 | 0.0 | 0.0 |
| Arg | 565 | Glu | 135 | NM | NM | 50.6 | 46.8 |
| Arg | 565 | Arg | 137 | NM | NM | 51.5 | 64.2 |
| Arg | 567 | Glu | 135 | NM | NM | 9.5 | 30.2 |
| Arg | 568 | Glu | 135 | NM | NM | 80.5 | 67.2 |
| Arg | 568 | Phe | 136 | NM | NM | 26.3 | 42.4 |
| Glu | 572 | Lys | 300 | NM | NM | 42.2 | 43.4 |
| Lys | 575 | Val | 298 | NM | NM | 22.0 | 31.1 |
| Arg | 576 | Val | 298 | NM | NM | 50.0 | 32.4 |
| Arg | 576 | Gly | 299 | NM | NM | 40.1 | 20.7 |
| Arg | 576 | Lys | 300 | NM | NM | 53.7 | 34.9 |
| Ala | 579 | Val | 298 | NM | NM | 37.1 | 38.3 |
| Interprotomer counterclockwise | | | | | | | |
| Val | 548 | Arg | 568 | 24.2 | 30.2 | 0.0 | 0.0 |
| Val | 548 | Asn | 571 | 23.1 | 48.6 | 0.0 | 0.0 |
| Val | 548 | Arg | 574 | 3.9 | 31.4 | 0.0 | 0.0 |
| Val | 548 | Lys | 575 | 9.4 | 31.7 | 0.0 | 0.0 |
| Ile | 549 | Ala | 564 | 13.9 | 32.7 | 0.0 | 0.0 |
| Leu | 551 | Leu | 578 | 1.1 | 28.6 | 0.0 | 0.0 |
| Glu | 552 | Arg | 565 | 39.3 | 20.2 | 0.0 | 0.0 |
| Glu | 552 | Arg | 567 | 44.1 | 58.0 | 0.0 | 0.0 |

| | | | | | | | |
|---|---|---|---|---|---|---|---|
| Glu | 552 | Arg | 568 | 69.8 | 39.4 | 0.0 | 0.0 |
| Glu | 552 | Arg | 574 | 3.3 | 33.5 | 0.0 | 0.0 |
| Tyr | 553 | Asn | 560 | 5.2 | 27.3 | 0.0 | 0.0 |
| Glu | 555 | Arg | 568 | 48.1 | 29.6 | 0.0 | 0.0 |
| Glu | 555 | Arg | 574 | 3.5 | 26.9 | 0.0 | 0.0 |
| Gln | 556 | Leu | 563 | 10.4 | 28.8 | 0.0 | 0.0 |
| Gln | 556 | Ala | 564 | 33.2 | 22.6 | 0.0 | 0.0 |
| Gln | 556 | Arg | 567 | 32.2 | 31.8 | 0.0 | 0.0 |
| Gln | 556 | Arg | 568 | 36.0 | 24.8 | 0.0 | 0.0 |
| Phe | 559 | Arg | 567 | 28.0 | 19.2 | 0.0 | 0.0 |
| Phe | 559 | Asn | 571 | 35.0 | 15.9 | 0.0 | 0.0 |

Contacts are quantified as the percentage of the accumulated simulation time in which any two non-hydrogen atoms from the two residues are within 4 Å of each other, and formed more than 25% of the simulation time, in simulations of either the inclined or flat orientations of the tail, with either 100 mM or 300 mM $K^+$. Results are sorted according to the protomer of the residue being contacted, and then by the residue number in the C-terminal tail forming the contacts. Rows are colored according to the segment being contacted: loop2a (residues 120-130, light green), loop2b (residues 131-137, dark green), loop4 (gray), loop6 (blue), TM12 (yellow) and the C-tail (white). The numbering of the loops indicates the TM segment that it follows, thus loop2 connects TM2 and TM3. NM indicates contacts not measured due to their location in the poorly sampled region following A564 toward the end of the C-terminal tail.

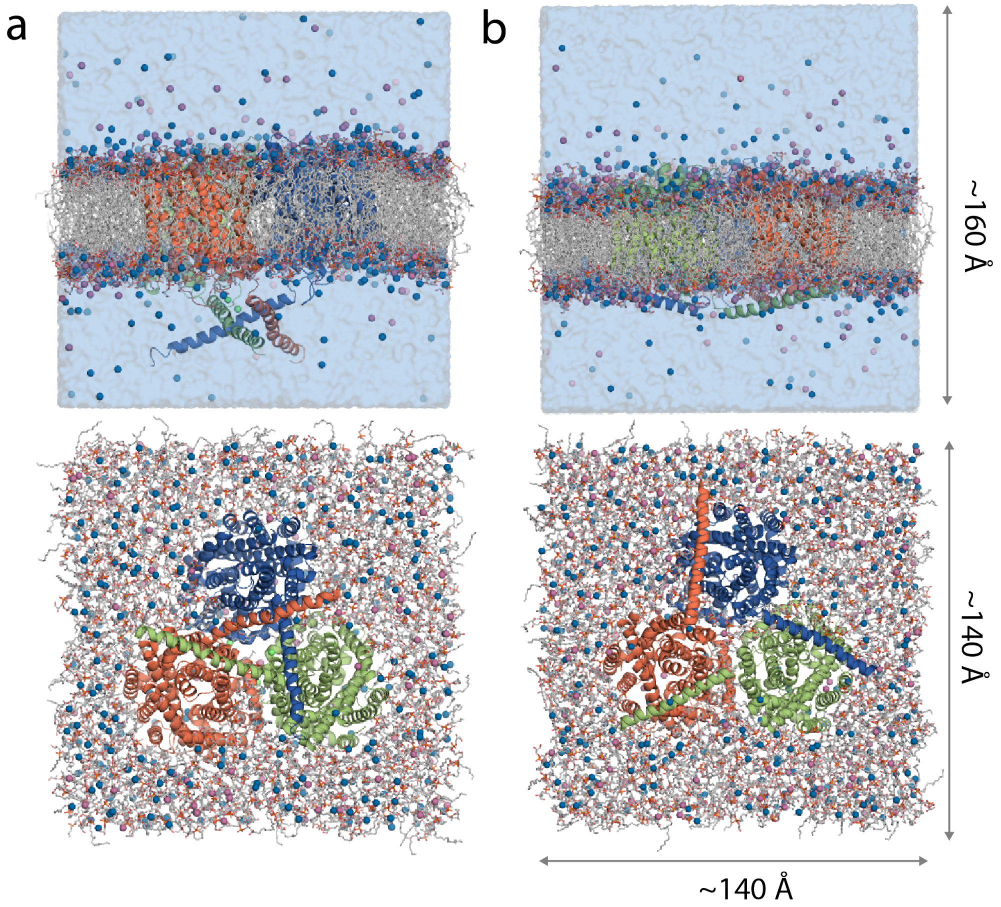


**Figure S1. Trimers of BetP embedded in a hydrated POPG lipid bilayer**. The C-terminal tails are either **(a)** inclined away from the transporter core, as found in X-ray crystal structures, or **(b)** lay flat along the membrane. In both cases, these tails interact with segments in neighboring protomers, as indicated by blue, orange, and green coloring. However, the differing orientations of the helices result in contacts with different regions of the protein. The protein (cartoon helices) is viewed from the plane of the membrane (top panels) or from the cytoplasm (lower panels), with POPG molecules shown as sticks, and ions shown as spheres for sodium (blue), potassium (purple) and chloride (green). Water is shown in the top panels as a blue surface and omitted from the lower panel for clarity.

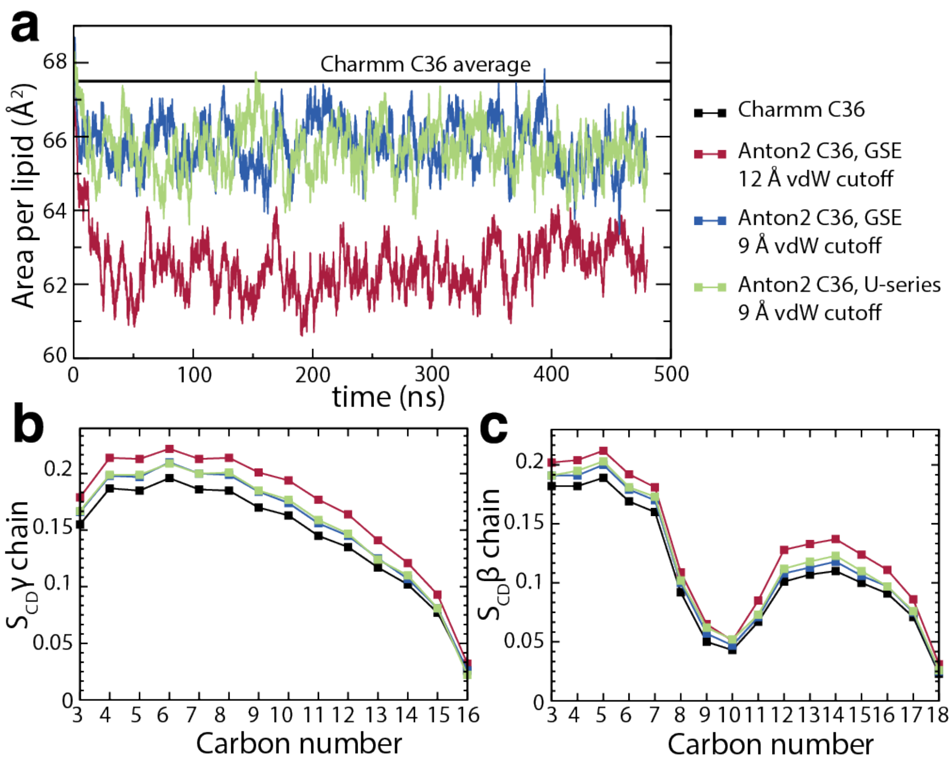


**Figure S2. Properties of a hydrated POPG lipid bilayer simulated using Charmm or Anton2. (a)** The area per lipid computed for three Anton2 simulations using van der Waals distance cut-offs of 12 Å (red) or 9 Å (blue, green), and either the GSE (red, blue) or U-series (green) treatment of the long-range electrostatics. The average value for the simulation performed using the Charmm package (published in [2]) is shown (black line). **(b, c)** Deuterium order parameters for the γ **(b)** and β chains **(c)** of the POPG lipids, colored according to panel a.

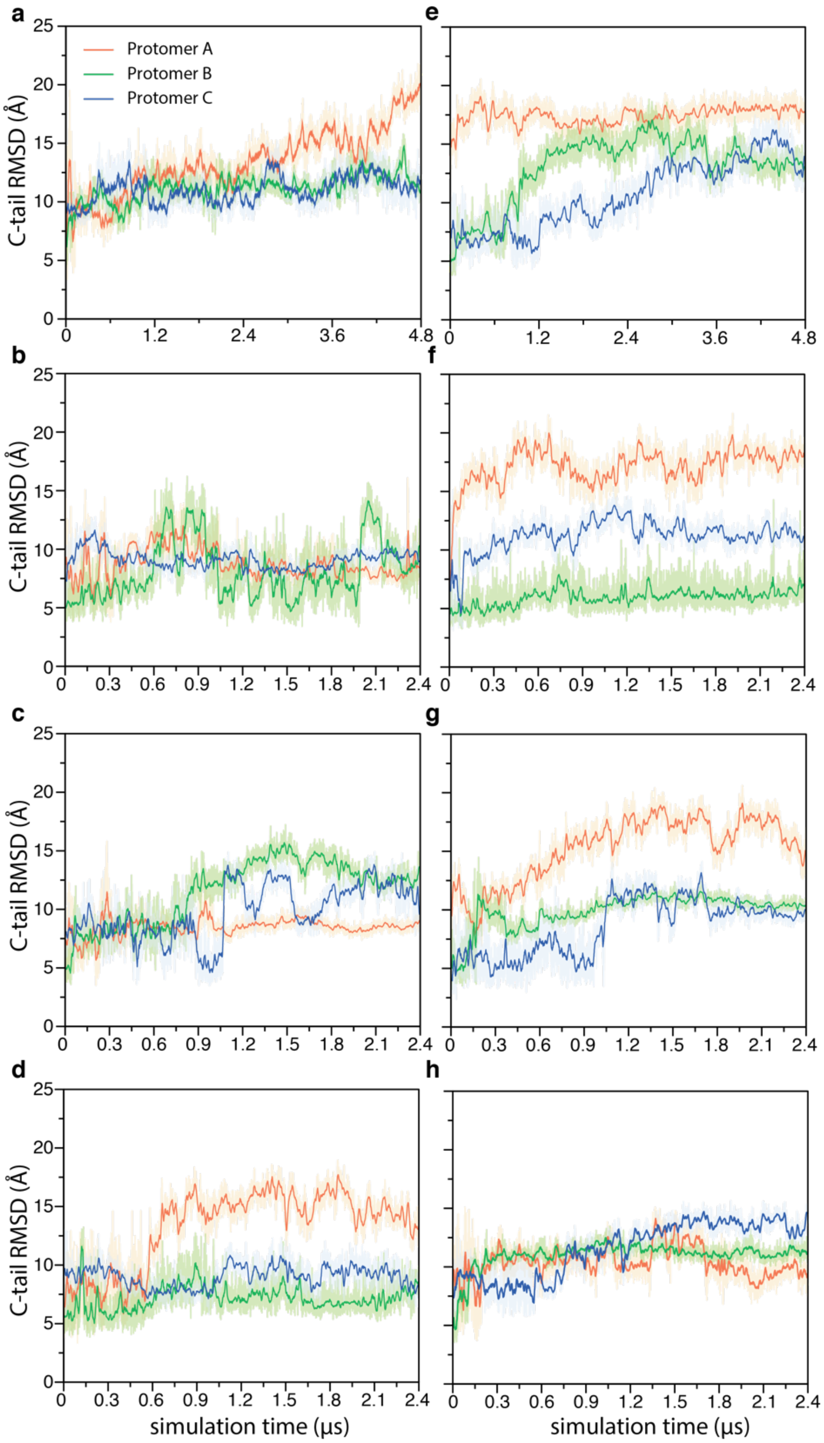


**Figure S3. Stability of the C-terminal tails derived from the crystal structure (inclined) in the complete set of simulations.** RMSD of each C-terminal tail after aligning the trimer using the transmembrane regions. The reference structure is the initial structure. **(a-d)** Results for four trajectories carried out using 100 mM $K^+$. **(e-h)** Results for four trajectories carried out using 300 mM $K^+$. Panels a and b are also shown in Figure 1.

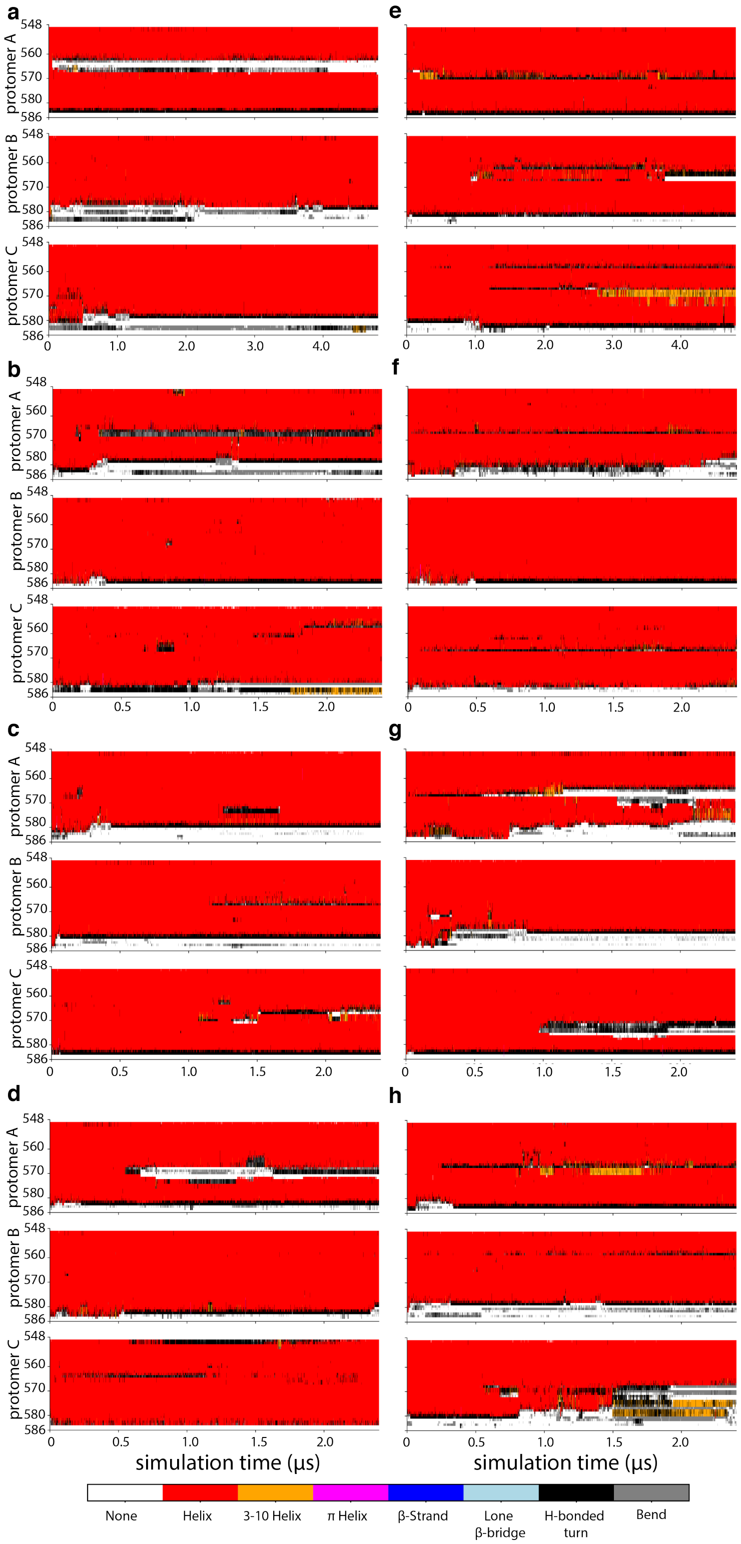


**Figure S4. Secondary structure of the C-terminal tail derived from the crystal structure (inclined) in the complete set of simulations.** The secondary structure computed using DSSP, plotted for every residue in each C-terminal tail using the indicated color scheme. **(a-d)** Results for four trajectories carried out using 100 mM $K^+$. **(e-h)** Results for four trajectories carried out using 300 mM $K^+$. Panels a and b are also shown in Figure 1.

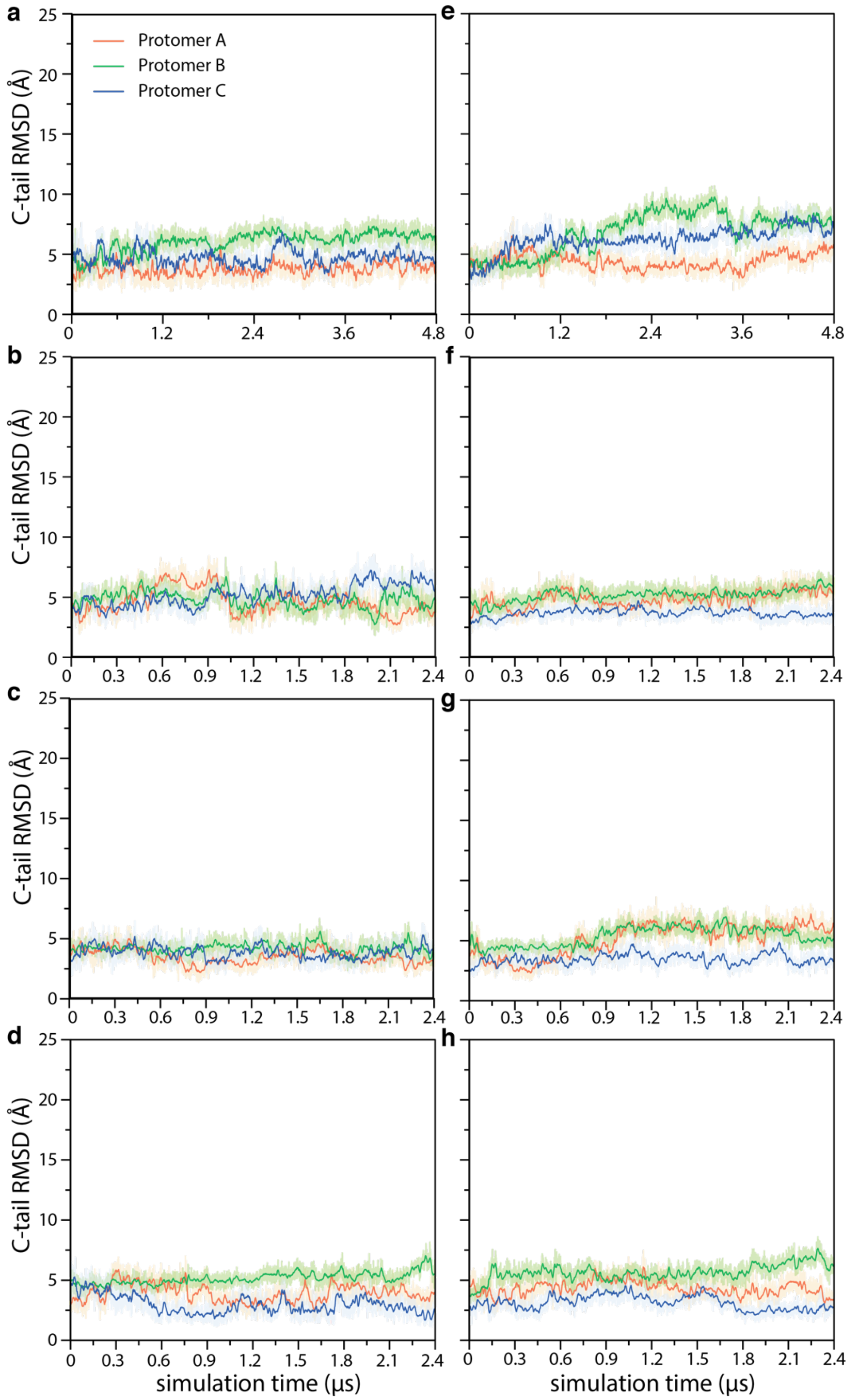


**Figure S5. Stability of the first half of the C-terminal tail in the simulations initiated from the crystal structure (inclined) in the complete set of simulations.** RMSD of residues 548 to 564 in the C-terminal tail of each protomer after aligning on the transmembrane regions. The reference structure is the initial structure. **(a-d)** Results for four trajectories carried out using 100 mM $K^+$. **(e-h)** Results for four trajectories carried out using 300 mM $K^+$.

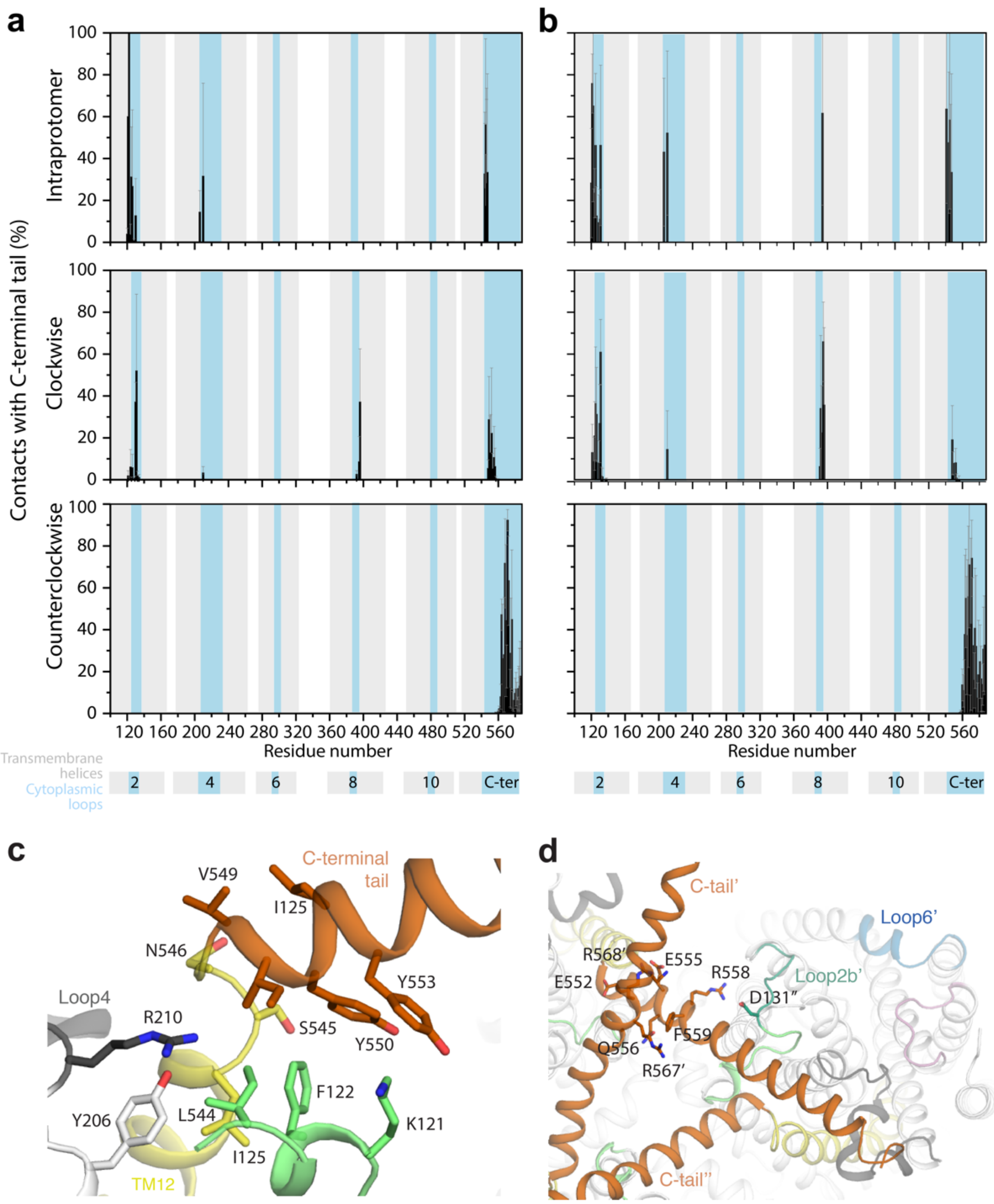


**Figure S6. Interactions between the first half of the C-terminal tail and the rest of the trimer in the simulations initiated from the crystal structure (inclined). (a,b)** Contacts computed for residues 548 to 564 of each C-terminal tail with the three adjacent protomers, i.e., intraprotomer, clockwise, or counterclockwise, assuming a view of the trimer from the cytoplasm. Contacts are defined as any two non-hydrogen atoms within 4.2 Å and plotted as bars for the mean (black) and standard deviation (error bars) over the three protomers and over all available simulations. Results are shown for the simulations of the BetP trimer performed with either 100 mM $K^+$ **(a)** or 300 mM $K^+$ **(b)**. The location of the cytoplasmic, periplasmic and TM regions of BetP is indicated by the legend. **(c, d)** Representative simulation frame, showing contacts of the C-terminal segment (orange) with the cytoplasmic regions of the same protomer (c) or with the adjacent protomers (d), in the clockwise ('') or counterclockwise directions ('). The protein is shown in cartoon representation colored yellow for TM12, light green for loop2a (residues 120-130), dark green for loop2b (residues 131-137), gray for loop4, and white for the remainder of the protein.

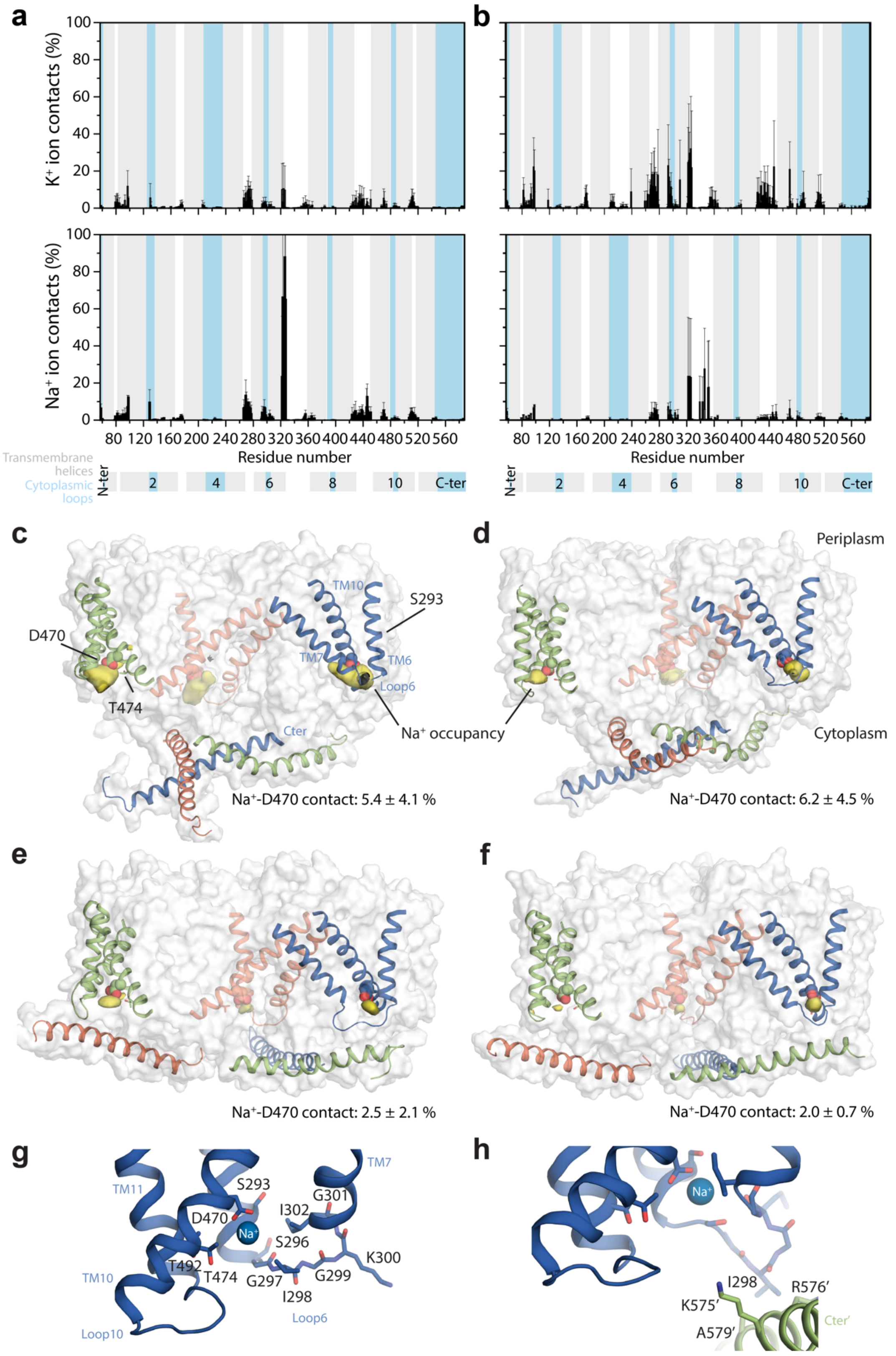


**Figure S7. Interactions between cations and the BetP trimer. (a, b)** Contacts formed during simulations initiated with the inclined orientation of the C-terminal tail, are defined as any ion ($K^+$, upper panels, or $Na^+$, lower panels) within 3.5 Å of any non-hydrogen atom of a residue in BetP. Contacts are plotted as an average (black bars) and standard deviation (error bars) across the three protomers and all available simulations. Results are shown for the simulations performed with either 100 mM $K^+$ **(a)** or 300 mM $K^+$ **(b)**. The location of the cytoplasmic (blue), periplasmic (white) and TM (gray) regions of BetP is indicated by the legend. **(c-f)** Sodium ion occupancy map (yellow surface), showing only densities close to the cytoplasmic surface and present in all three protomers, for clarity. The protein is shown in gray surface representation and as cartoon helices for TM6, TM7, TM10 and the C-terminal tail, colored by protomer (orange, blue and green). Asp470 is shown as spheres, and Ser293 is shown as sticks. Simulations were initiated with either **(c, d)** the inclined orientation of the C-terminal tails found in X-ray crystallographic structures, or **(e, f)** the flat orientation observed in EM maps. The bulk concentration of $K^+$ was either 100 mM (c, e) or 300 mM (d, f). Mean and standard deviation of the contact time between $Na^+$ ions and Asp470 are indicated. **(g, h)** Closeup of the sodium ion when coordinated by Asp470 in simulations in which the C-terminal tail is inclined (g), or in a flat orientation (h). Representative snapshots with high coordination numbers are shown with the ion (sphere), and nearby amino acids (sticks) from cytoplasmic ends of TM6, TM7, TM10, and TM11 (cartoon helices and sticks). In (h), the C-terminal tail of the adjacent protomer forms close contacts with residues in the TM6-7 loop. Individual residues are labelled. Segments from an adjacent protomer are labelled with prime (′).

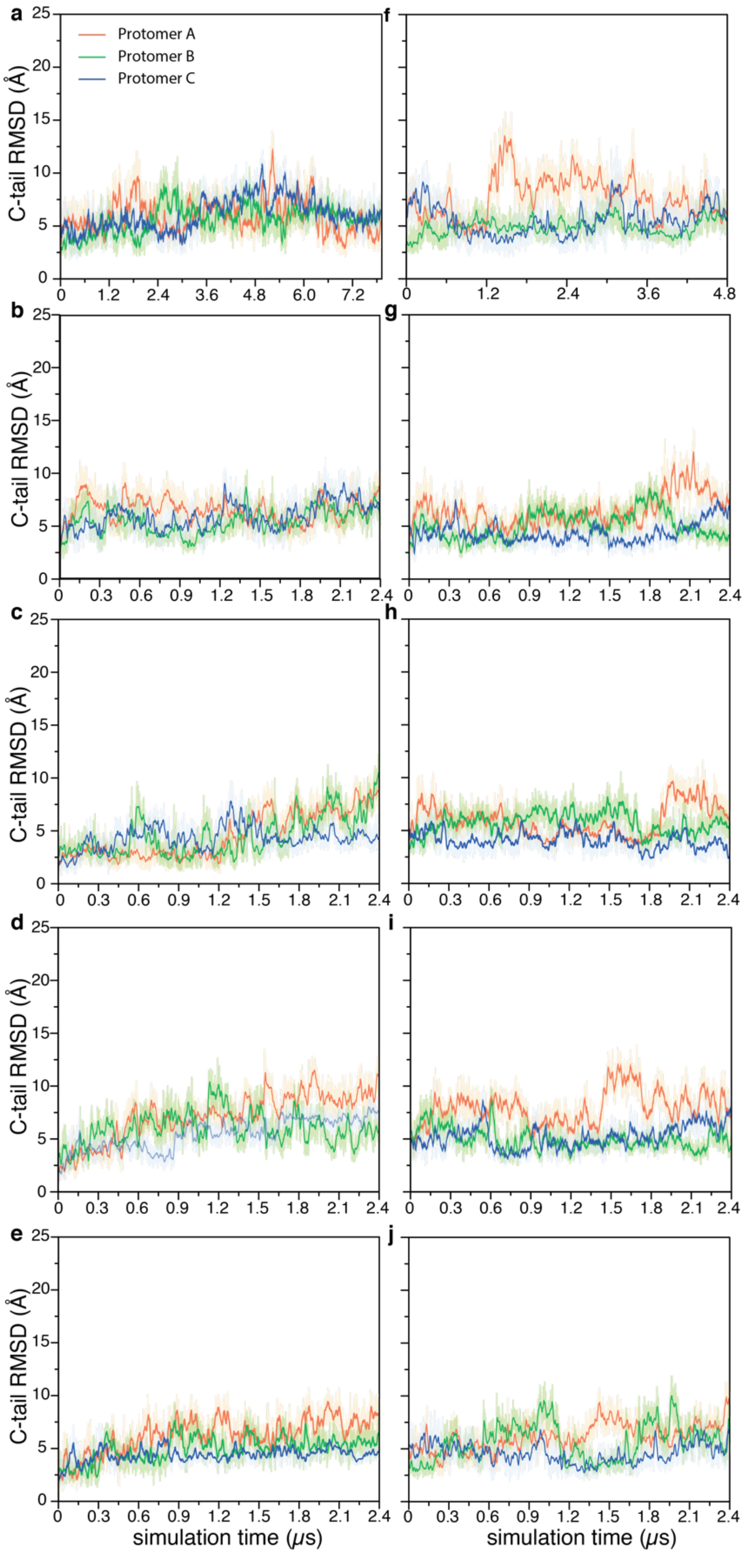


**Figure S8. Stability of the C-terminal tails derived from the EM data (flat) in the complete set of simulations of the BetP trimer.** RMSD of each C-terminal tail in BetP after aligning on the transmembrane regions. The reference structure is the initial structure. Simulations were carried out using **(a-e)** 100 mM $K^+$ or **(f-j)** 300 mM $K^+$. Panels a and f are also shown in Figure 3.

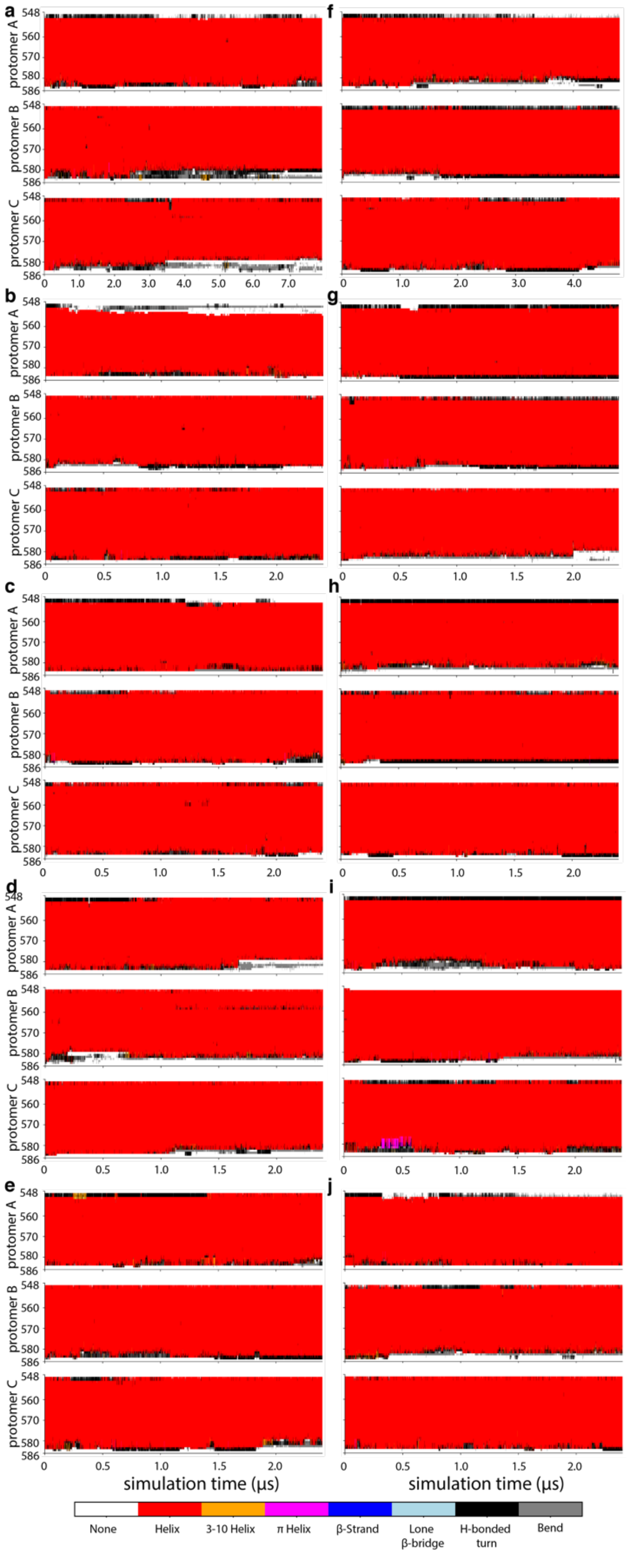


**Figure S9. Secondary structure of the C-terminal tails derived from the EM data (flat) in the complete set of simulations.** The secondary structure computed using DSSP, plotted for every residue in each C-terminal tail as a function of simulation time, using the indicated color scheme for trajectories using (**a-e)** 100 mM $K^+$ or **(f-j)** 300 mM $K^+$. Panels a and f are also shown in Figure 3.

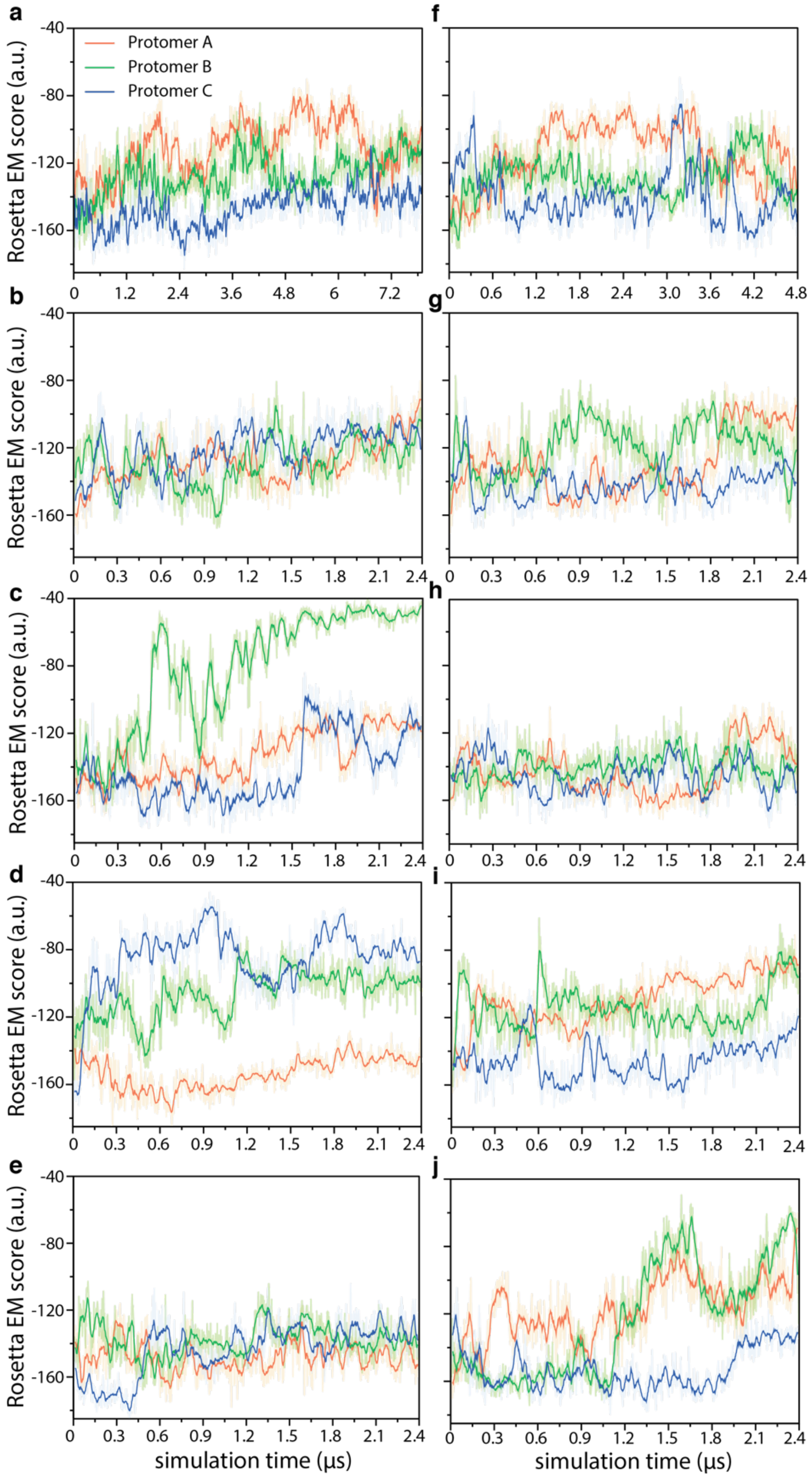


**Figure S10. Agreement of the C-terminal tail conformations in the complete set of "flat" simulations with the EM maps.** The Rosetta Electron Microscopy [5] score of each C-terminal tail plotted as a function of simulation time for five trajectories using **(a-e)** 100 mM $K^+$, or **(f-j)** 300 mM $K^+$.

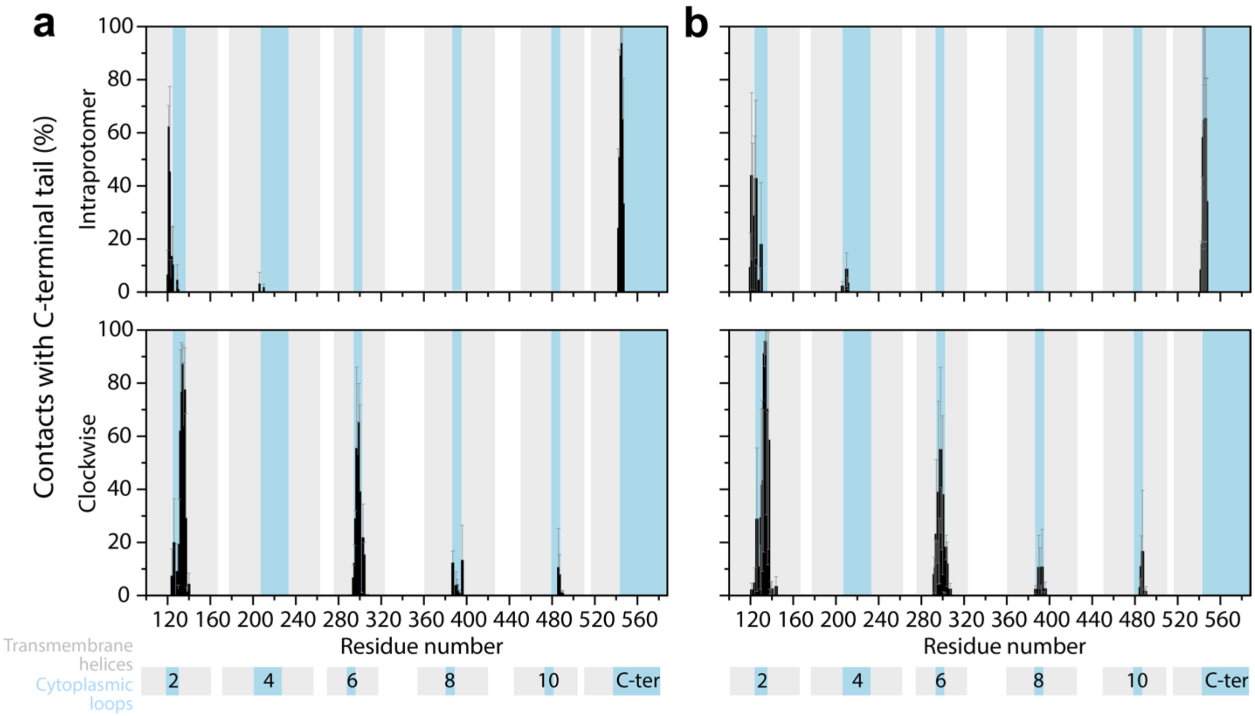


**Figure S11. Interactions between the C-terminal tail and the rest of the trimer in simulations derived from the EM data (flat).** Contacts were computed for each C-terminal tail with two adjacent protomers, either intraprotomer or clockwise, assuming the trimer is viewed from the cytoplasm. Counterclockwise contacts are not reported because none were detected in any snapshots of our simulations. Contacts are defined as any two non-hydrogen atoms within 4.2 Å and are plotted as bars for the mean (black) and standard deviation (error bars) over all available simulations. Results are shown for the simulations performed with either 100 mM $K^+$ **(a)** or 300 mM $K^+$ **(b)**. The location of the cytoplasmic (blue), periplasmic (white) and TM (gray) regions of BetP is indicated by the legend.

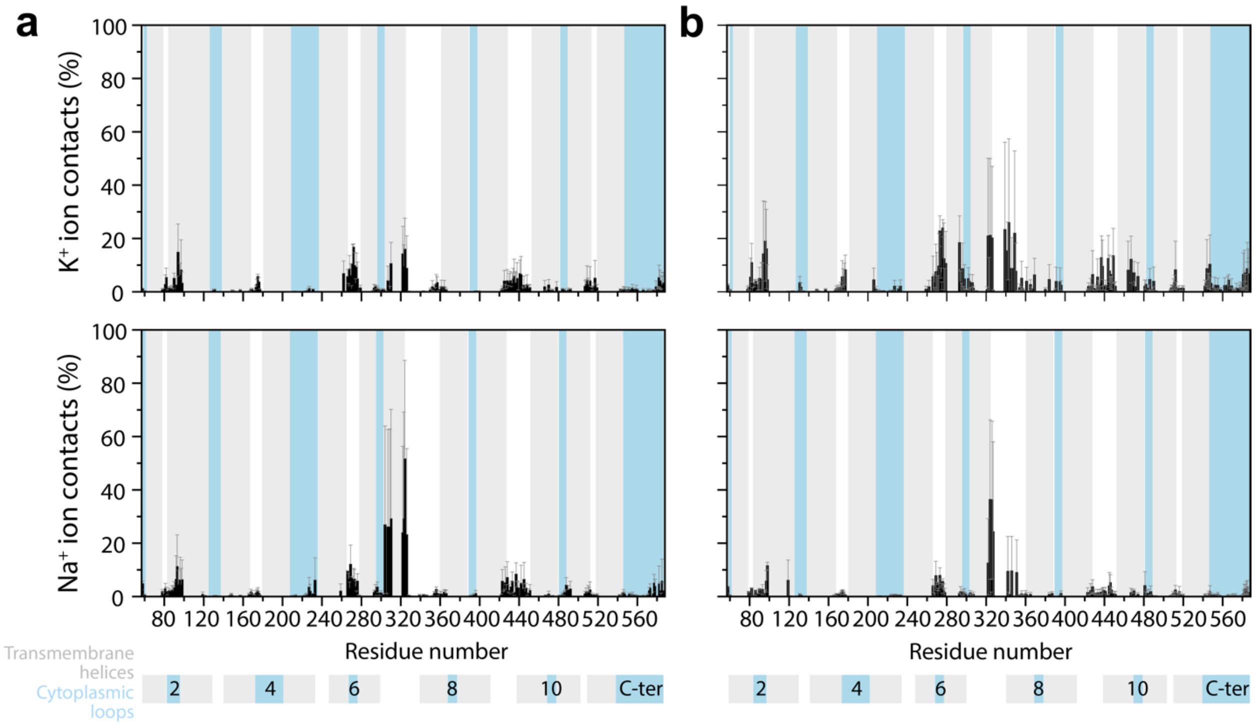


**Figure S12. Interactions between cations and the BetP trimer in simulations initiated from the EM data (flat).** Contacts are defined as any ion ($K^+$, upper panels or $Na^+$, lower panels) within 3.5 Å of any non-hydrogen atom of a residue in BetP. Contacts are shown as an average (black bars) and standard deviation (error bars) across the three protomers and all available simulations. Results are shown for the simulations performed with either 100 mM $K^+$ **(a)** or 300 mM $K^+$ **(b)**. The location of the cytoplasmic (blue), periplasmic (white) and TM (gray) regions of BetP is indicated by the legend.

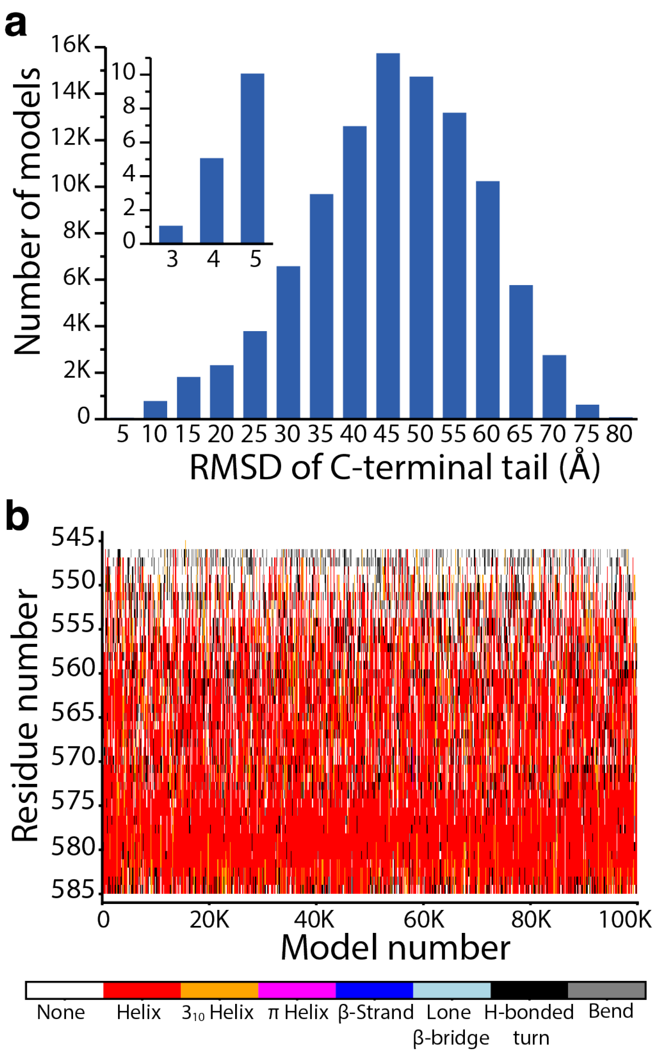


**Figure S13. Structural accuracy of *de novo* predictions of the BetP C-terminal tail. (a)** RMSD with respect to the EM structure (all-flat) for all 100,000 *de novo* models of the C-terminal tail, after aligning on the transmembrane regions. Inset: Close-up of the data for models with RMSD values < 5 Å. **(b)** The secondary structure of all 100,000 *de novo* models, computed using DSSP, plotted for every residue using the indicated color scheme.

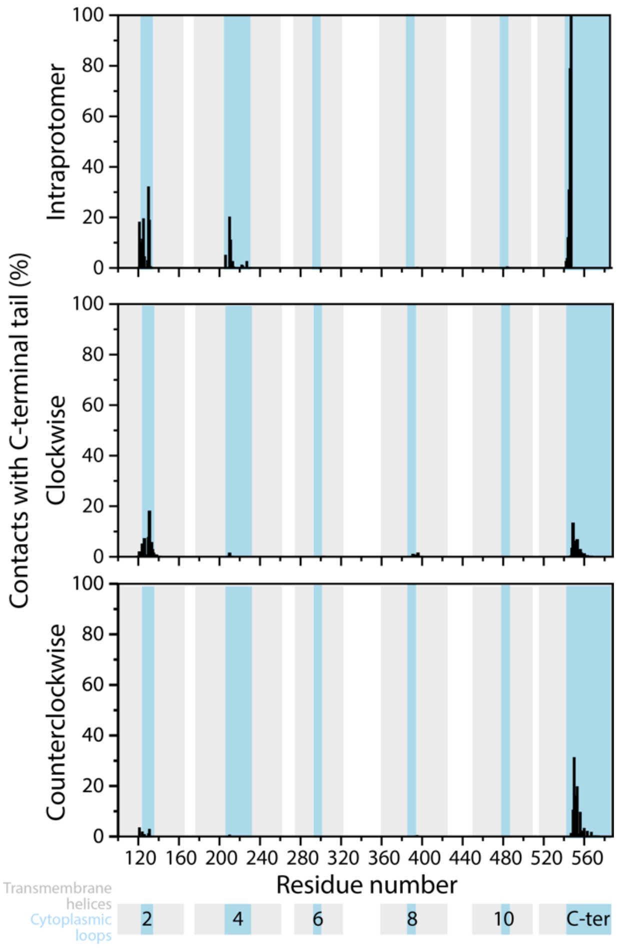


**Figure S14. Predicted interactions between *de novo* models of the C-terminal tail and the rest of the BetP trimer.** Contacts of the modeled C-terminal tail with individual residues in each of the three protomers of BetP, either the same protomer (top), the clockwise protomer (middle), or counterclockwise (lower), assuming a view of the trimer from the cytoplasm. Contacts are defined as any two non-hydrogen atoms within 4.2 Å. The location of the cytoplasmic (blue), periplasmic (white) and TM (gray) regions of BetP is indicated by the legend.

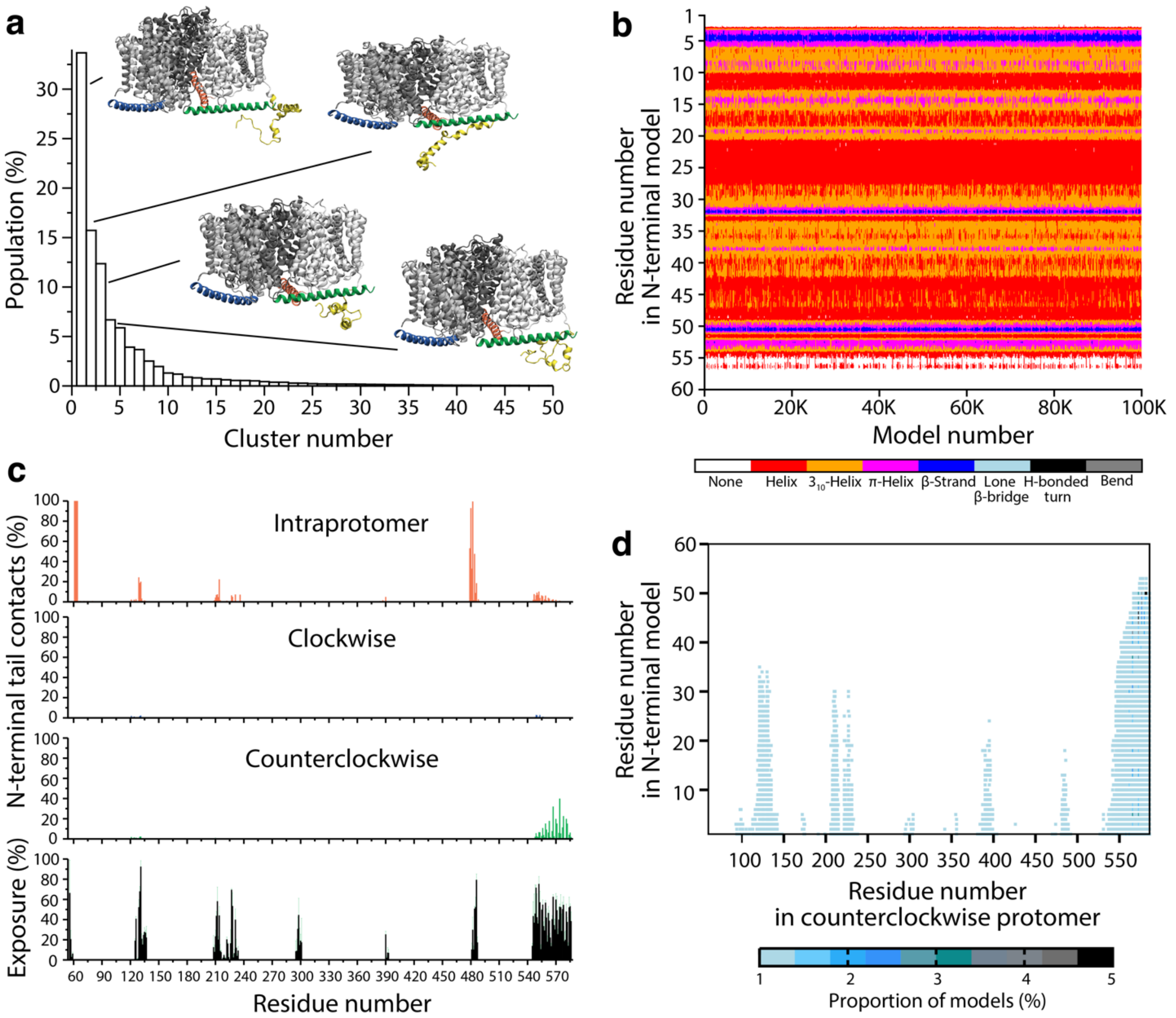


**Figure S15. Structural features and contacts formed by the *de novo* predicted N-terminal tail of BetP, when including residues 1 through 60. (a)** Clusters of 100,000 models of residues 1 to 60 of the N-terminal tail, based on their C-$\alpha$ RMSD. For the four most-populated clusters, the structure at the center of that cluster is shown as cartoon helices, viewed from the plane of the membrane. The transmembrane segments are colored three shades of gray, the C-terminal segments are orange, blue and green, and the predicted N-terminal segment is colored yellow. **(b)** The secondary structure in all 100,000 models of residues 1-60 of the N-terminal tail built considering residues 1 to 60, computed using DSSP and colored according to the legend. **(c)** Contacts of the modeled N-terminal tail residues 1 to 60 with individual residues in each of the three protomers of BetP, either intraprotomer (red bars), clockwise (blue bars), or counterclockwise (green bars), assuming a view of the trimer from the cytoplasm. Contacts are defined as any two non-hydrogen atoms within 4.2 Å. Bottom panel: Available contact surface in the cytoplasmic regions of the BetP trimer, excluding the N-terminal segment. Data is plotted as the mean (black) and standard deviation (green error bars) over the three protomers of the solvent-accessible surface area, relative to the maximum possible for each amino acid type in a GXG tripeptide. **(d)** Contacts between any residue of the modeled N-terminal tail (residues 1 to 60) and residues in the counterclockwise protomer. Points are colored according to the number of models in which that contact was observed as indicated by the legend.

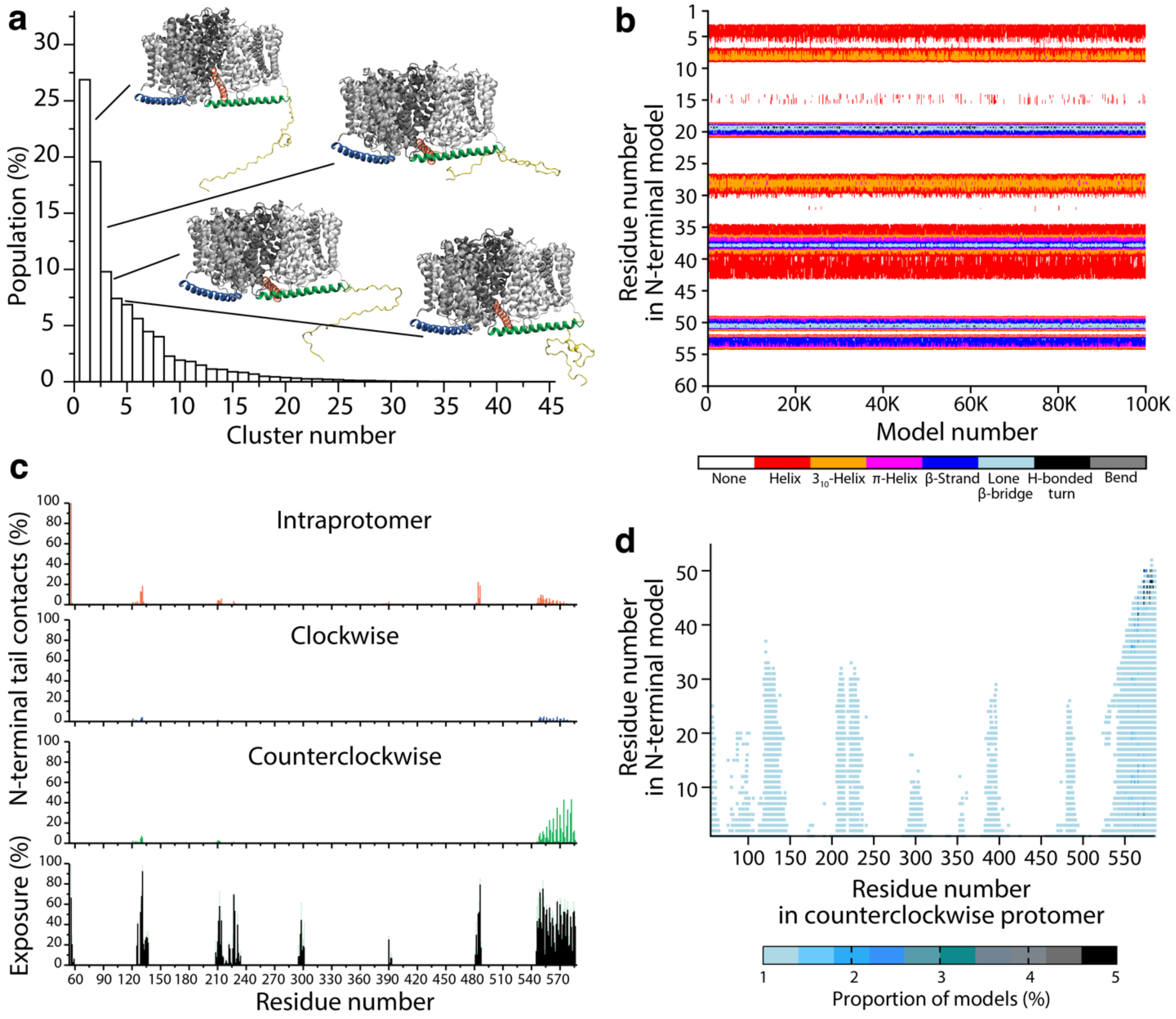


**Figure S16. Structural features of, and contacts formed by the *de novo* predicted N-terminal tail of BetP excluding information from local structural fragments. (a)** Clusters of 100,000 models of residues 1 to 55 of the N-terminal tail, based on their Cα RMSD. No local fragment information was included during model building. For the four most-populated clusters, the structure at the center of that cluster is shown as cartoon helices, viewed from the plane of the membrane. The transmembrane segments are colored three shades of gray, the C-terminal segments are orange, blue and green, and the predicted N-terminal segment is colored yellow. **(b)** The secondary structure in all 100,000 models of residues 1-55 of the N-terminal tail, computed using DSSP and colored according to the legend. **(c)** Contacts of the modeled N-terminal tail (residues 1 to 55) with individual residues in each of the three protomers of BetP, either intraprotomer (red bars), clockwise (blue bars), or counterclockwise (green bars), assuming a view of the trimer from the cytoplasm. Contacts are defined as any two non-hydrogen atoms within 4.2 Å. Bottom panel: Available contact surface in the cytoplasmic regions of the BetP trimer, excluding the N-terminal segment. Data is plotted as the mean (black) and standard deviation (green error bars) over the three protomers of the solvent-accessible surface area, relative to the maximum possible for each amino acid type in a GXG tripeptide. **(d)** Contacts between any residue of the modeled N-terminal tail (residues 1 to 60) and residues in the protomer located counterclockwise to the one containing the modelled N-terminal tail. Points are colored according to the number of models in which that contact was observed, as indicated by the legend.

**Supplementary Materials References**

1. Brooks BR, Brooks CL, Mackerell AD, Nilsson L, Petrella RJ, Roux B, et al. CHARMM: The Biomolecular Simulation Program. J Comput Chem. 2009;30(10):1545-614
2. Venable RM, Brown FLH, Pastor RW. Mechanical properties of lipid bilayers from molecular dynamics simulation. Chem Phys Lipids. 2015;192:60-74
3. Shan YB, Klepeis JL, Eastwood MP, Dror RO, Shaw DE. Gaussian split Ewald: A fast Ewald mesh method for molecular simulation. J Chem Phys. 2005;122(5)
4. Predescu C, Lerer AK, Lippert RA, Towles B, Grossman JP, Dirks RM, et al. The u-series: A separable decomposition for electrostatics computation with improved accuracy. J Chem Phys. 2020;152(8)
5. DiMaio F, Tyka MD, Baker ML, Chiu W, Baker D. Refinement of Protein Structures into Low-Resolution Density Maps Using Rosetta. J Mol Biol. 2009;392(1):181-90
